# Determinants of centromeric nucleosome conformation

**DOI:** 10.64898/2026.05.15.724335

**Authors:** Emeleeta A. Paintsil, Joshua Lee, Sarah S. Meidl Zahorodny, Jeff Wereszczynski, Emma A. Morrison

## Abstract

Centromeric chromatin is defined by the presence of the histone H3 variant CENP-A, which forms a specialized nucleosome required for kinetochore assembly. Compared to canonical H3 nucleosomes, CENP-A nucleosomes exhibit an open DNA conformation that leaves an additional 13 base pairs of DNA accessible at the entry and exit sites. While the CENP-A αN-helix has previously been implicated in promoting this enhanced DNA breathing, the contributions of the intrinsically disordered N-terminal tail and adjacent latch regions of CENP-A to nucleosome conformation remain unknown. The intrinsically disordered N-terminal regions of histone H3 are known to facilitate interactions with DNA to stabilize overall nucleosome conformation. Here, we utilized a combination of MNase digestion assays and coarse-grained molecular dynamics simulations of nucleosomes containing targeted swaps of the H3 and CENP-A tail, latch, and αN-helix regions to systematically test the contribution of each N-terminal histone region to nucleosomal DNA conformation. Substitution of individual H3 N-terminal regions with those of CENP-A opened the DNA at the entry/exit sites and increased DNA accessibility. While any single CENP-A N-terminal region was sufficient to open the canonical nucleosomal DNA conformation, replacement of any single CENP-A N-terminal region with its H3 counterpart was insufficient to restore the closed DNA conformation characteristic of canonical H3 nucleosomes. Instead, progressive incorporation of multiple H3-derived regions produced increasingly closed DNA conformations, demonstrating that the H3 tail, latch, and αN-helices act cooperatively to stabilize wrapped nucleosomal DNA. Taken together, these findings demonstrate that the more restricted DNA breathing of canonical nucleosomes arises from coordinated contributions across multiple N-terminal regions and suggest that the multi-region redundancy in the conformational flexibility of the centromeric nucleosome could emphasize the importance of retaining flexibility in the centromeric nucleosome, even upon post-translational modification and binding to structural proteins.

**SIGNIFICANCE:** The centromere is marked by nucleosomes containing CENP-A, which adopt a more open and accessible DNA conformation than canonical nucleosomes. However, the molecular determinants underlying this difference remain unclear. Previous structural investigations of the centromeric nucleosome have placed less emphasis on the intrinsically disordered N-terminal regions of CENP-A. Here, we systematically dissect the contributions of the N-terminal tail, latch, and αN-helix via MNase digestion assays and molecular dynamics simulations on nucleosomes containing chimeras of H3 and CENP-A. We demonstrate that no individual H3-derived region is sufficient to impart a closed conformation to the nucleosomal DNA. Instead, multiple regions act together to stabilize DNA, revealing that nucleosome conformation is controlled by concerted histone-DNA interactions.

## INTRODUCTION

Chromatin regulation begins at the level of its basic subunit, the nucleosome. The canonical nucleosome is comprised of a histone octamer core containing two copies each of the H3, H4, H2A, and H2B histones wrapped by ∼147 base pairs (bp) of DNA (1). Each of the canonical histones has a similar secondary structure, with a histone fold comprised of three primary alpha helices (α1, α2, and α3) along with histone-fold extensions (1). The histones have disordered “tail” regions at the N- (and C- for H2A) termini that protrude from the nucleosome core and interact with DNA in a dynamic conformational ensemble (2–4). These histone tails are also important sites of interaction with chromatin-associating proteins (5). A major mechanism of chromatin regulation is through incorporation of histone variants, which are non-canonical histone sequences that serve specialized regulatory functions within nucleosomes and chromatin (6, 7). These sequence variations can function to differentially regulate chromatin via both direct and indirect mechanisms by (i) directly altering the stability, dynamics, and/or conformations of the nucleosome, (ii) introducing or removing binding sites for regulatory or structural proteins, and (iii) introducing or removing sites of histone post-translational modifications (PTMs). Histone variants are assembled into nucleosomes depending on factors such as cell cycle phase, chromatin type and location, and cell type (6, 7). Disruptions in the incorporation patterns of histone variants are associated with genomic instability and disease states, such as cancer, that arise from dysregulation at the chromatin level (8).

The centromere is a specialized structure of chromatin, marked by nucleosomes assembled with the H3 histone variant Centromeric Protein-A (CENP-A) (9). CENP-A is essential for proper chromosome segregation (10, 11). Dysregulation in the form of aberrant localization outside of the centromere leads to global disruption of chromosome structure (12–15) and is associated with cancer and poor prognosis (13, 16–19). CENP-A nucleosomes serve as the foundation of the constitutive centromere-associated network (CCAN), which is a complex of proteins that bridges the centromere to the kinetochore (20, 21). Several segments of the CENP-A sequence that diverge from canonical H3 are known to be important for indirect mechanisms of chromatin regulation, specifically by introducing binding sites for CCAN proteins: the RG loop for CENP-N binding (22) and the C-terminus to bind CENP-C (23).

Molecularly, the CENP-A nucleosome structure differs from that of the H3 nucleosome: the centromeric nucleosome has a more open and flexible DNA conformation. The X-ray crystal structure of the CENP-A nucleosome lacks DNA density for ∼13 bp of the entry and exit DNA (24), and cryo-electron microscopy (cryo-EM) further confirmed the increased heterogeneity and flexibility (25–27). Moreover, atomic force microscopy experiments also captured the flexible nature of the CENP-A nucleosome (28–30). In combination with these structural studies, biochemical assays assessing the susceptibility of DNA ends to MNase and exonuclease digestion support differences between H3 and CENP-A nucleosomal DNA accessibility (24, 26, 31–33). This observation holds within the context of nucleosome arrays, suggesting this feature is relevant in chromatin systems too (34). An interesting dichotomy exists for centromeric nucleosomes: although the centromeric nucleosome is more open and flexible at DNA entry/exit sites and more elastic than the canonical nucleosome (25, 28, 35, 36), nucleosome arrays with CENP-A are more condensed than with H3 (34). We speculate that this apparent contradiction in nucleosome properties plays an important role in variant-specific function, and an understanding of the molecular-level details of nucleosome conformation may provide insight into these contrasting properties (37).

The CENP-A sequence differs substantially from H3, more so than any other H3 variant: H3 and CENP-A share ∼49% sequence identity across the entire histone. There are two intrinsically disordered N-terminal regions: i) the tail and ii) the residues that pass through the gyre of DNA and flank the αN-helix, which we refer to here as the histone “latch” region [the term latch is adapted from (38), but we use it here to refer to a shorter fragment that excludes residues in the αN-helix] **(Figure 1A)**. CENP-A and H3 share ∼33% and 24% sequence identity in the latch and tails, respectively. These regions are known to facilitate interactions with nucleosomal DNA that are integral to the stability and conformation of canonical nucleosomes (39). Canonical and variant H3 sequences also possess a short helix, termed the αN-helix, between the N-terminal disordered regions and the core histone fold. The αN-helices of CENP-A and H3 are ∼54% identical in sequence.

**Figure 1.**
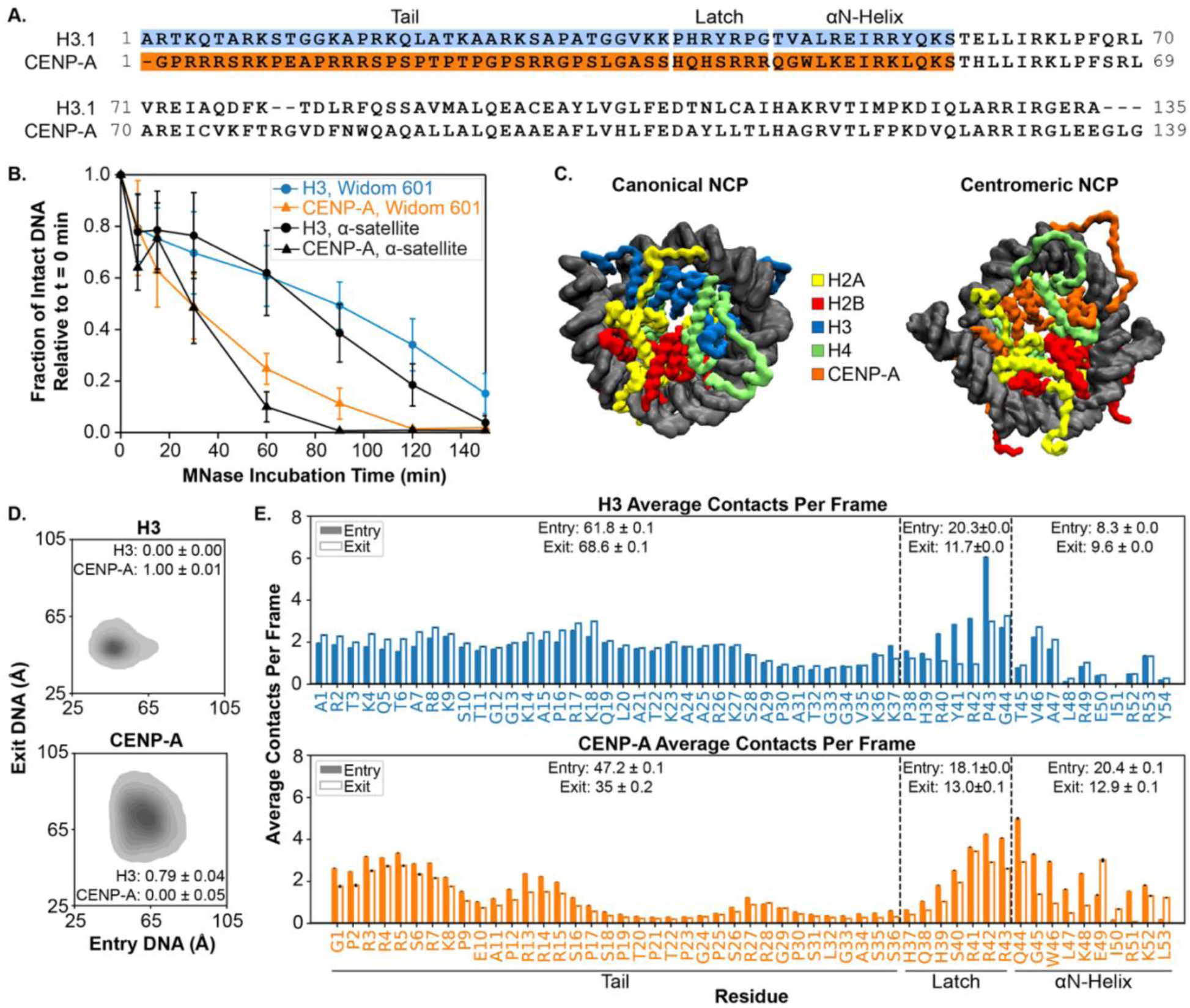
Wild-type NCPs display distinct DNA conformations and histone-DNA interaction profiles. **A.** Alignment of the H3 and CENP-A histone sequences with N-terminal regions labeled**. B.** MNase digestion time course for NCPs reconstituted with H3 (circles) or CENP-A (triangles) and Widom 601 (blue, orange) or α-satellite (black) DNA. Plots represent the average of a minimum of n=3 assays, and error bars represent the standard deviation of these data. **C.** Representative snapshot from CG simulations of H3 and CENP-A NCPs (H2A: yellow, H2B: red, H3: blue, H4: green, CENP-A: orange) with α-satellite DNA. **D.** Plots of end-to-dyad distances from CG simulations of H3 and CENP-A NCPs with α-satellite DNA. KL divergence values normalized on a scale from 0 to 1 are shown to indicate the similarity to both H3 and CENP-A, wherein 0 indicates complete identity to that system, and 1 indicates maximal computed divergence. Errors represent the standard deviation for KL. **E.** Per-residue bar plots of average contacts with DNA for the tail, latch, and αN-helix regions for H3 (top; blue) and CENP-A (bottom; orange) from CG simulations of H3 and CENP-A NCPs with α-satellite DNA. Contacts are shown for entry (solid bars) and exit (open bars) tails. Per-residue values represent the mean of n=4 independent simulation replicates, with error bars representing the standard error of the mean across replicates. Total average contacts for each region are above the data bars, with errors representing propagated SEM calculated from the individual per-residue SEM values.

The more open and flexible DNA conformation of the centromeric as compared to canonical nucleosomes has been attributed to the shortened αN-helix of the CENP-A histone (25). The canonical H3 αN-helix is defined by residues T44-S57, but in crystal and cryo-EM structures with CENP-A, only residues L47-S56 are observed as helical; residues Q44-W46 of CENP-A are disordered (24, 26). Despite the reported importance of the αN-helix in determining nucleosome accessibility, other regions of the histone may contribute to nucleosome conformation. For example, the H3 variant H3.Y also imparts increased accessibility to nucleosomal DNA, although to a lesser extent than CENP-A (31). The αN-helices of H3.1 and H3.Y have the same length and differ only at two residues, and the difference in accessibility between H3.Y and canonical H3 was instead attributed to disruptions to interactions with the DNA ends that are mediated by latch and core residues (31). In the context of the canonical nucleosome, deletion of the histone tails increases DNA accessibility (40–42). However, the impact of the tails on the centromeric nucleosome is unknown. Although it is assumed that the H3 and CENP-A tails have similar roles within the nucleosome, an investigation of the relative contributions from the CENP-A tail on DNA conformation has yet to be conducted. We ask whether the N-terminal regions of CENP-A play a role in maintaining the unique DNA conformation of CENP-A NCPs.

In this study, we utilize a combination of MNase digestion assays and coarse-grained molecular dynamics (MD) simulations to systematically investigate the contributions of the N-terminal regions of H3 and CENP-A to overall nucleosome conformation. While the CENP-A tail, latch, and αN-helix each promote a more accessible conformation within the canonical nucleosome, the entire N-terminal region of H3 is needed for a closed NCP conformation in the centromeric nucleosome. Through our MD simulations, we observe that contacts between the H3 and CENP-A N-terminal regions and DNA are influenced by the adjacent sequence context and contribute to DNA conformation. Here, we identify a role for the histone tail and latch region in maintaining the DNA conformation of the H3 and CENP-A NCPs. These findings provide new insight into how coordinated interactions between intrinsically disordered histone regions contribute to nucleosome conformation, and we speculate that they contribute to the specialized structural dynamics required for proper centromeric chromatin function.

## MATERIALS AND METHODS

### Purification of canonical histones via inclusion bodies

H3.1 (UniProt P68431), H4 (UniProt P62805), H2A (UniProt P0C0S8), and H2B(UniProt P62807) histones were expressed and purified as previously described (43). Briefly, histones were purified from inclusion bodies and purified in 7 M Urea buffer [7 M urea, 100 mM NaCl, 20 mM sodium acetate, pH 5.2, 5 mM β-mercaptoethanol 1 mM EDTA] over Q-Sepharose Fast Flow resin (GE Healthcare) and SP-Sepharose Fast Flow resin (GE Healthcare) or Proteindex IEX CM Agarose 6 Fast Flow resin (Marvelgent Biosciences). Histones were dialyzed against ddH20 with 1 mM β-mercaptoethanol, lyophilized, and stored at −20°C until use in NCP reconstitutions. Histone purity was confirmed by ESI mass spectrometry (Finnigan LTQ, Thermo Electron Corporation, San Jose, CA). The H3 with CENP-A *a*N and H3 with CENP-A latch chimera histones were also expressed and purified via this method. All other H3 and CENP-A chimera histones were purified following the CENP-A/H4 tetramer purification method.

### CENP-A/H4 tetramer expression and purification

CENP-A (UniProt P49450) was expressed and purified as a tetramer with H4 (UniProt P62805) using a bicistronic plasmid system (the pST39 vector) (44–46). All subsequent CENP-A/H3 chimera histones, except the H3 with CENP-A αN and H3 with CENP-A latch, were made from this plasmid backbone. Briefly, H4/CENP-A tetramer was expressed in BL21(DE3) or T7 Express LysY competent *E. coli* (New England BioLabs) and grown in 2xYT media (RPI) at 37°C, induced with 0.2 mM IPTG, and incubated at 18°C for 16 hours. Cells were lysed and sonicated in buffer [20 mM NaPO4, 750 mM NaCl, 1 mM EDTA, 1 mM Benzamidine-HCl, 10 mM β-mercaptoethanol], pH 7, with additives of 0.25 mg/mL DNase I, 0.5 mg/mL lysozyme, 0.5% Triton X-100. Lysates were clarified by centrifugation at 257,300 RCF for 30 minutes and filtered (0.22 µm) before FPLC (BioRad) purification. Soluble tetramer was purified over HiTrap SP Fast Flow 5mL Column (Cytiva) via FPLC (BioRad), dialyzed against 4 L of buffer [15 mM NaPO_4_, 30 mM MOPS, 1 M NaCl, 10 ppm CaCl_2_, 10 mM β-mercaptoethanol, pH 6.8], and then purified over Foresight™ CHT™ Type II Ceramic Hydroxyapatite Column (BioRad).

The boundaries for the N-terminal regions used in designing the chimera sequences are defined as follows (**Figure S1**): the tail is residues 1-37 in H3 and 1-36 in CENP-A; the latch is residues 38-44 in H3 and 37-43 in CENP-A; the *a*N-helix is residues 45-54 in H3 and 44-53 in CENP-A; and the core is residues 55-135 in H3 and 54-139 in CENP-A.

### Preparation of DNA

147bp Widom 601 DNA and 147bp α-satellite DNA were prepared for reconstitution as previously described (47, 48). A plasmid containing 32 repeats of 147-bp Widom 601 sequence (ATCGAGAATC CCGGTGCCGA GGCCGCTCAA TTGGTCGTAG ACAGCTCTAG CACCGCTTAA ACGCACGTAC GCGCTGTCCC CCGCGTTTTA ACCGCCAAGG GGATTACTCC CTAGTCTCCA GGCACGTGTC AGATATATAC ATCCGAT) or a 147-bp α-satellite DNA sequence (ATCAAATATC CACCTGCAGA TTCTACCAAA AGTGTATTTG GAAACTGCTC CATCAAAAGG CATGTTCAGC TCTGTGAGTG AAACTCCATC ATCACAAAGA ATATTCTGAG AATGCTTCCG TTTGCCTTTT ATATGAACTT CCTCGAT) (45) was transformed into NEB 5-alpha competent *E. coli* (NEB - C2987I) and grown in 2xYT media pH 7.5. Plasmids were purified using an alkaline lysis method followed by phenol-chloroform extraction. The plasmid was digested by EcoRV-HF enzyme (NEB - R3195L), and DNA fragments were purified from the vector using polyethylene glycol (PEG) precipitation and HiTrap DEAE Fast Flow Column (GE Healthcare). Lastly, 147-bp DNA underwent ethanol precipitation at −20°C. Prior to reconstitution, DNA was resuspended in high salt buffer [20 mM Tris-HCl, pH 7.5, 2M KCl, 1 mM EDTA, 0.5 mM benzamidine, 1 mM DTT] to a final concentration of 10 µM DNA.

### NCP reconstitution

Canonical and centromeric nucleosomes were reconstituted similarly to as previously described (43, 45, 48). Refolded H3/H4 or purified soluble CENP-A/H4 tetramers were combined with H2A/H2B dimer and 10 µM 601 DNA in high salt buffer [20 mM Tris-HCl, pH 7.5, 2 M KCl, 1 mM EDTA, 0.5 mM benzamidine, 1 mM DTT]. NCPs were reconstituted by dialyzing over a slow desalting gradient into buffer without salt [20 mM Tris-HCl, pH 7.5, 1 mM EDTA, 0.5 mM benzamidine, 1 mM DTT]. Canonical NCPs were reconstituted in a final 1:1:2 molar ratio of DNA: Tetramer: Dimer, wild-type CENP-A and H3/CENP-A chimera NCPs were reconstituted in a 1:1:1.8 molar ratio. NCPs were concentrated and heat-shocked at 55°C for an hour. NCPs were purified over a 10%-40% sucrose gradient (BioComp Instruments). NCP purity was determined by assessing the absorbance at 260 nm of the sucrose gradient profile and 5% Native-PAGE. NCP histone integrity was determined by 18% SDS-PAGE (**Figure S2 and S3**).

**MNase assays.** MNase assays were carried out at 171 nM NCP in MNase reaction buffer [42 mM Tris-HCl (pH 7.5), 15 mM NaCl, 2.5 mM CaCl_2_, 1.8 mM DTT], similar to published conditions (31), and digested with 6.75 units of Micrococcal Nuclease (NEB, M0247S) per 100 µL of reaction. Reactions were incubated at 37°C. The reaction time was started upon addition of MNase. Samples were collected from the bulk reactions at each time point (7, 15, 30, 60, 90, 120, and 150 min) and immediately quenched by diluting the sample 1:1 into Novex™ TBE-Urea Sample Buffer (2X) (Thermo Fisher, LC6876), heating at 95°C for 10 mins, and then immediately storing on ice prior to loading on a gel. MNase samples were separated on 10% TBE-Urea Criterion gels (BioRad, 3450089) run in 1X TBE running buffer (diluted from BioRad 1610733), at a constant voltage of 220 V for 65 min at room temperature. Prior to running samples, gels were pre-run at a constant voltage of 220 V for 45-60 min at room temperature. MNase digestion gels were quantified using Image Lab software (BioRad) and plotted with IgorPro 9 software (Wavemetrics). Briefly, gel lanes and DNA bands were added manually. Band boundaries were adjusted by eye using the *Lane Profile* with a background subtraction of 50.6 mm (disk size). The *Adjusted Volume Intensity* data values were obtained for the intact DNA at each time point. Within each time series, the data were scaled to 1 with respect to the 0 min incubation *Adjusted Volume Intensity*. The resulting scaled values were plotted as a function of MNase incubation time.

### Coarse-grained nucleosome construction and simulation setup

Coarse-grained (CG) versions of all atomistic systems were made with version 3.1 of CafeMol (49). The first post-equilibration frame of a single replica per system was used as the starting frame for all CG simulations. All full-length CG NCPs were constructed with alpha-satellite DNA. CG systems used the AICG2+ protein and 3SPN2.C DNA potential models (50, 51). Each amino acid was simplified into a single Cα bead, whereas DNA residues were simplified into three beads representing the base, sugar, and phosphate. The base beads were placed at the N1 and N3 atoms for purines and pyrimidines, respectively. The sugar bead was placed at the center of mass of the sugar ring, and the phosphate bead represents the phosphate atom in the DNA backbone. DNA and protein chains interacted through excluded volume, Debye-Hückel electrostatics, native contacts, and hydrogen-bonded interactions carried over from the all-atom starting PDB (52). Standard integer charges were used for protein residues. A default scaling factor of 1.0 was used to keep the nucleosomes stable throughout the trajectories at an ionic concentration of 0.150 M. A temperature of 300 K was used, with temperature maintained via Langevin dynamics. The friction coefficient, dielectric constant, and Debye length at 0.150 M were 0.843 and 78.0, and approximately 8 Å, respectively. In the Debye-Hückel treatment used here, salt concentrations are incorporated via bulk electrostatic screening, lacking any explicitly modeled ions (53). Consequently, effects such as divalent-ion interactions, local ion concentrations, and other spatially heterogeneous ionic environments are not captured (54). Therefore, the specified salt concentrations should only be regarded as effective values chosen to maintain consistency rather than as direct representations of the local ionic environment. Electrostatic interactions were computed with a 20 Å cutoff distance. Phosphate bead charges were set to −0.6, considering the Oosawa Manning counterion condensation around the DNA (55, 56). A phosphate charge of −1.0 for DNA-protein interactions was used based on prior coarse-grained studies. Timesteps for all CG simulations were 0.3 Cafemol time units, which roughly corresponds to 30 ps of physical time (57). Similar to all-atom systems, all CG systems were simulated in replicas of four for 100,000,000 steps each, or the rough equivalent of 3 ms effective nucleosome kinetics time per replica (**Table S2**). Trajectories were saved every 1,000 steps. Convergence of CG simulations was assessed based on the end-to-dyad distributions used to characterize nucleosome opening. For each independent simulation, end-to-dyad distributions calculated using progressively larger fractions (10-90%) of the production trajectory were compared with the corresponding full-trajectory distribution using Jensen-Shannon divergence **(Figure S4)** (58).

### Simulation analyses

Root mean square deviation (RMSD) and root mean square fluctuation (RMSF), and end-to-dyad distance analyses were conducted using CPPTRAJ from AmberTools on post-equilibration data (59–61). RMSD and RMSF calculations were conducted after alignment to the Cα atoms of the alpha helices in the first frame H4 histone pair heavy atoms for consistency, due to all NCPs sharing the same H4 sequence despite differing in H3/CENP-A composition. End-to-dyad distances were calculated as the distance between the center of mass of the five base pairs of each DNA terminus and the dyad, which corresponds to base pair 74. Kullback-Leibler (KL) divergence (62, 63) calculations based on end-to-dyad data were performed for all NCPs, one calculation using H3 as a basis and a second with CENP-A as a basis. All KL values were normalized from 0 to 1 using min-max scaling with the entire CG end-to-dyad dataset. Hydrogen bond analyses between the N-terminal tails and DNA were performed in the built-in VMD tcl scripting terminal using in-house scripts (64). A donor-acceptor cutoff of 3.5 Å and a cutoff angle of 30° between D-H-A atoms were used. All intramolecular and DNA base pairing hydrogen bonds were excluded in the calculation.

Contact analyses between the protein and DNA were conducted using the MDAnalysis package (65, 66). Contacts were defined as any heavy atoms within 4 Å of one another between the two residue selections for all-atom systems and 8 Å for CG systems. Residues 1-37 in H3 and 1-36 in CENP-A were considered as the tails in all relevant nucleosomes. A latch region for H3 was considered residues 38-44 in H3 and 37-43 in CENP-A. Residues 45-54 in H3 and 44-53 in CENP-A were defined as the αN-helices. In spiral contact analyses, contacts between histone tails and DNA were computed in each trajectory frame and assigned to individual DNA bases. Each contacted base was further classified as “inside” or “outside” based on the sign of the cosine angle (greater than zero inside, less than zero outside) between a local nucleotide orientation vector (C1’ atom to C3’ atom for all-atom and base to sugar atoms for CG) and the vector from the DNA base position to the histone core center of mass. Contact frequencies were averaged over independent replicas to obtain mean values and standard errors of the mean (SEMs).

All analyses were plotted using the matplotlib (67) and/or seaborn (68) packages within python. Unless otherwise specified, reported error bars for all required analyses represent the SEM and treat all four replicates per system as independent runs. For analyses reporting SEM across replicates, quantities were first averaged over the trajectory of each independently initialized and equilibrated simulation, and SEM was subsequently calculated across the four replicate-level averages. Individual trajectory frames were not treated as independent statistical observations. Both all-atom and CG data were analyzed using similar scripts for shared analyses.

## RESULTS

### Coarse-grained simulations recapitulate biochemical determinations of differential DNA accessibility between canonical and centromeric nucleosomes

We began our study by establishing a baseline for H3 and CENP-A nucleosomal DNA accessibility. We digested NCPs assembled with two different DNA sequences: one set with the artificial Widom 601 DNA sequence (69) and the other with an α-satellite DNA sequence intended to mimic native centromeric DNA (70, 71) **(Figure 1B, Figures S5A, S5B, S6)**. We observe that the Widom 601 and α-satellite H3 NCPs had a normalized band volume of 0.6 ± 0.1 and 0.6 ± 0.2 after 60 minutes, respectively, which is greater than the values of 0.25 ± 0.06 and 0.09 ± 0.05 obtained for Widom 601 and α-satellite CENP-A NCPs, respectively **(Figure 1B)**. This demonstrates that the DNA in the CENP-A NCP is more accessible than in the H3 NCP, regardless of DNA sequence, which is consistent with previous experimental results (26, 31, 33). Because trends hold between DNA sequences, we conducted the remainder of our MNase digestion assays using NCPs reconstituted with the Widom 601 sequence. We note that while MNase assays are commonly used to investigate the accessibility of nucleosomal DNA ends (26, 31, 33), and we interpret our results in these terms, we cannot definitively exclude contributions from DNA gaping or other forms of conformational distortions that could increase DNA accessibility to the MNase. Nevertheless, a side-by-side comparison between MNase and Exonuclease III assays with CENP-A and H3.3 NCPs reveals (qualitatively) high similarity between the results (33), supporting that exonuclease activity is the dominant MNase activity being observed in these assays.

We turned to molecular dynamics (MD) simulations of the canonical H3 and CENP-A nucleosomes to gain insight into the specific intermolecular interactions that dictate nucleosomal conformation. We initially conducted all-atom MD simulations, but these sampled relatively short timescales and showed few changes in DNA conformation (**Supplemental Methods**, **Supplemental Results**, **Figures S7-S20**, **Tables S1-S13**). We instead focused our efforts on coarse-grained (CG) simulations of the canonical H3 and centromeric NCPs using α-satellite DNA to test whether we could observe flexible DNA opening in the CENP-A nucleosome on longer timescales. CG simulations were performed in quadruplicate for 3 ms of effective nucleosome kinetics time (57), with the first 0.6 ms dedicated to equilibration **(Figure 1C)**. To gain insight into differences in DNA conformation between CG NCPs, we analyzed the end-to-dyad distances of both the entry and exit DNA termini from the dyad base pair. We observed that the H3 NCP adopts a closed DNA conformation at both entry and exit sites while CENP-A adopts a more open DNA conformation **(Figure 1D)**.

To identify which residues might be driving the differing DNA conformations, we computed the per-residue contacts between the N-terminal regions of each histone and DNA from our CG simulations (**Figure 1E**). We observe that H3 has consistent DNA contacts along much of the intrinsically disordered N-terminal region, with each residue averaging 1-2 contacts per frame. In contrast, the CENP-A N-terminal region primarily displays DNA contacts peaking where clusters of arginine residues are present (e.g., arginine residues 3-5, 13-15, 27-28, and 41-43 show more contacts relative to other CENP-A residues). When we divide the contacts into sections, we note that the H3 tail has more total contacts (entry/exit: 61.8/68.6) compared to the CENP-A tail (entry/exit: 47.2/35.0). In the latch region, the number of DNA contacts is similar between CENP-A (entry/exit:18.1/13.0) and H3 (entry/exit: 20.3/11.7), suggesting a similar extent of DNA-latch interactions between variants. In contrast, there is a noticeable difference in the total DNA-αN-helix contacts between the H3 and CENP-A NCPs: the H3 αN-helix has roughly half as many total contacts (entry/exit:8.3/9.6) as the CENP-A αN-helix (entry/exit: 20.4/12.9).

Taken together, these observations suggest that the combination of increased tail contacts and decreased αN-helix contacts in the H3 NCP may promote a closed DNA conformation, while the decreased tail contacts and increased αN-helix contacts in the CENP-A NCP may underlie the open DNA conformation. This led us to hypothesize that the differential DNA interactions across the three N-terminal regions (the tail, latch, and αN-helix) collectively maintain the respective conformations of the H3 and CENP-A nucleosomes.

### Full N-terminal histone swaps through the αN-helix largely determine nucleosomal DNA conformation

After establishing that centromeric NCPs exhibit greater DNA accessibility than canonical NCPs and identifying differential patterns of DNA contacts across the N-terminal regions of CENP-A and H3, we sought to determine if the N-terminal region of the histone dictates NCP DNA conformation, or whether the rest of the histone core (i.e., all regions C-terminal to the αN-helix) contributes. To test this, we designed chimera histones to swap the full N-terminal region (i.e., tail, latch, and αN-helix) between H3 and CENP-A.

We probed the DNA conformation of the chimera-histone NCPs via MNase assay and CG simulations. MNase digestion of H3_Tail+Latch+αN_CENP-A_Core_ NCP displayed more restricted DNA accessibility than CENP-A NCP, on par with H3 NCP **(Figure 2A)**. Consistent with this, the end-to-dyad analysis of CG simulations shows that the H3_Tail+Latch+αN_CENP-A_Core_ NCP exhibits a dominant closed DNA state (**Figure 2B).** On the other hand, the CENP-A_TailLatch+αN_H3_Core_ NCP was more readily digested by MNase, although not quite to the extent of CENP-A NCP (**Figure 2A),** and has a primarily open DNA state in CG simulations, with some prominent closed DNA conformations as well (**Figure 2B).** Data from these two chimeras suggest that the combined effects of all N-terminal regions (tail, latch, αN-helix) are largely sufficient for maintaining the closed and open DNA conformations of canonical and centromeric NCPs, respectively.

**Figure 2.**
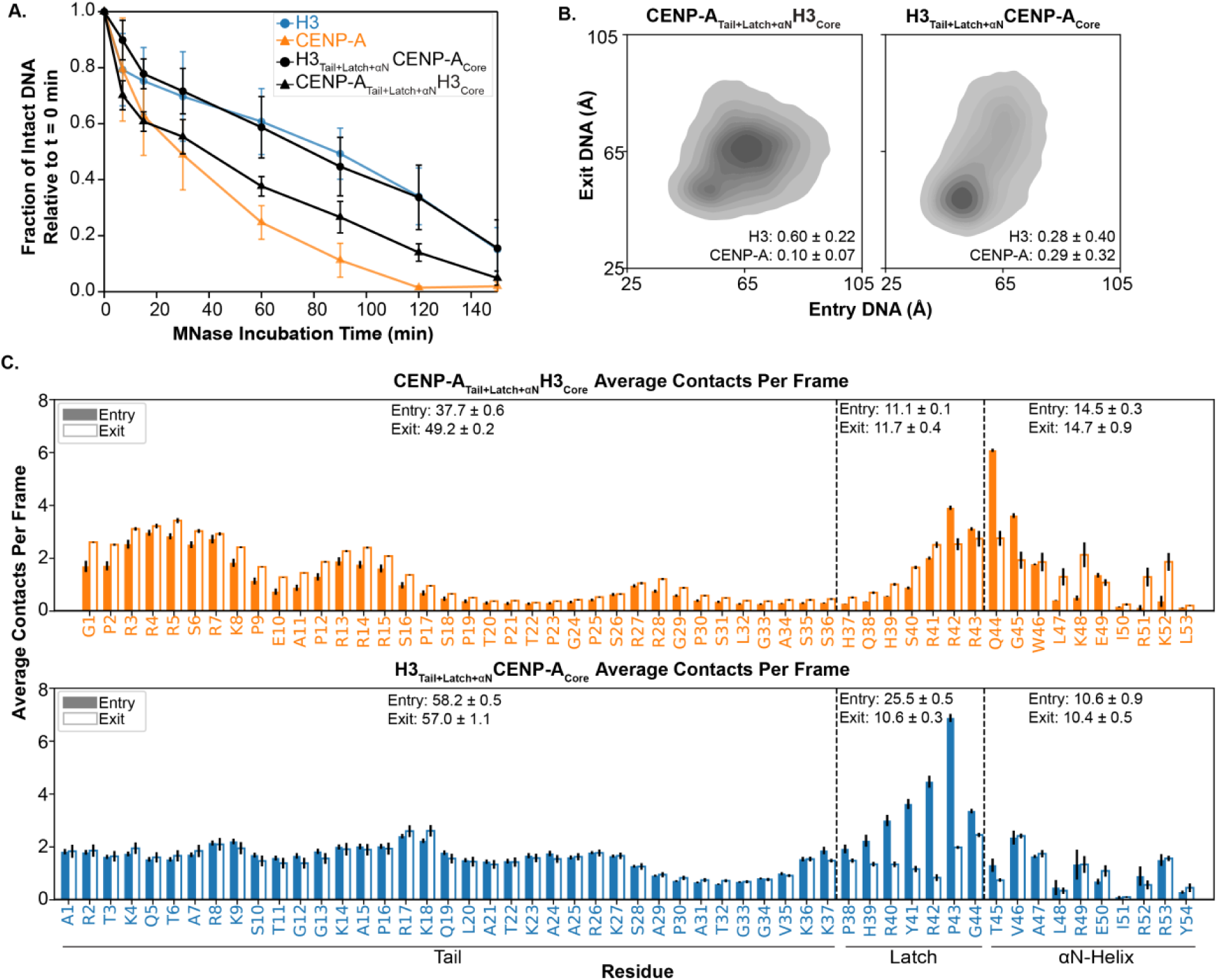
Chimeras swapping the tail through αN-helix show that the N-terminal histone-fold extension largely determines nucleosomal DNA conformation. **A** MNase digestion plot of H3_Tail+Latch+αN_CENP-A_Core_ (black circles) and CENP-A_Tail+Latch+αN_H3_Core_ (black triangles) with H3 (blue circles) and CENP-A (orange triangles) for comparison. Plots represent the average of a minimum of n=3 assays, and error bars represent the standard deviation of these data. NCPs use Widom 601 DNA. **B.** Plots of end-to-dyad distances from CG simulations of H3_Tail+Latch+αN_CENP-A_Core_ and CENP-A_Tail+Latch+αN_H3_Core_ NCPs with α-satellite DNA. Plots are inset with KL divergence values, with errors representing standard deviation. **C.** Per-residue bar plots of average contacts with DNA for the tail, latch, and αN-helix regions for H3_Tail+Latch+αN_CENP-A_Core_ and CENP-A_Tail+Latch+αN_H3_Core_ from CG simulations of NCPs with α-satellite DNA. Contacts are shown for entry (solid bars) and exit (open bars) tails. Blue bars and text represent amino acid sequence originating from H3; orange bars and text represent amino acid sequence originating from CENP-A. Per-residue values represent the mean of n=4 independent simulation replicates, with error bars representing the standard error of the mean across replicates. Total average contacts for each region are above the data bars, with errors representing propagated SEM calculated from the individual per-residue SEM values.

To quantify these trends in terms of similarity/divergence from both H3 and CENP-A wild-type (WT) NCPs, the end-to-dyad probability densities from CG simulations were used to conduct Kullback-Leibler (KL) divergence calculations (62, 63) against both H3 and CENP-A **(Table S9).** After normalization of KL values on a scale of 0-1, we find that the CENP-A_Tail+Latch+αN_H3_Core_ NCP exhibits a KL divergence of 0.60 ± 0.22 relative to the H3 reference, indicating a moderate deviation from H3-like behavior. The relatively large standard deviation suggests substantial variability across replicas, consistent with a heterogeneous conformational ensemble. It also has a KL divergence of 0.10 ± 0.07 relative to CENP-A NCP, suggesting that the DNA conformation is more similar to the CENP-A NCP than the H3 NCP. On the other hand, the H3_Tail+Latch+αN_CENP-A_Core_ has KL divergence values of 0.28 ± 0.40 relative to H3 and 0.29 ± 0.32 relative to CENP-A. This indicates an intermediate state with no strong bias toward either reference. These relatively large standard deviations again suggest substantial conformational heterogeneity across replicas.

Per-residue tail-DNA contacts along these N-terminal regions show that contacts are largely preserved from WT NCPs **(Figure 2C).** In H3_Tail+Latch+αN_CENP-A_Core_ NCP, H3 tails maintain broadly distributed and persistent contacts along the DNA, with a moderate decrease in total exit tail contacts. Although the total number of DNA contacts remains similar in CENP-A_Tail+Latch+αN_H3_Core_ NCP as compared to WT CENP-A NCP, the CENP-A tail flips to having more contacts in the exit tail than the entry tail [entry/exit: 37.7/49.2 vs. 47.2/35.0 (WT)]. The latch and αN-helix of H3_Tail+Latch+αN_CENP-A_Core_ show a slight increase in total DNA contacts relative to WT H3 [latch entry/exit: 25.5/10.6 vs. 20.3/11.7 (WT); αN entry/exit: 10.6/10.4 vs. 8.3/9.6] while the total DNA contacts in the latch and αN-helix of CENP-A_Tail+Latch+αN_H3_Core_ decrease compared to WT CENP-A [latch entry/exit: 11.1/11.7 vs. 18.1/13 (WT); αN entry/exit: 14.5/14.7 vs. 20.4/12.9]. This suggests that the DNA-contact behaviors of the histone N-terminal regions are moderately affected by the sequence of the main globular histone fold region.

Together, these data demonstrate the importance of the N-terminal region of the H3 variant (i.e., H3 vs. CENP-A) in dictating the nucleosomal DNA conformation. In H3 systems, the number of contacts between histone tail residues and DNA is relatively consistent throughout the tail, whereas in CENP-A systems, the contacts are more localized to arginine clusters within the tails, specifically near the N-termini and latch regions. These more localized contacts may create disparate binding modes between the systems, as the CENP-A tails may more strongly anchor to the DNA at the N-termini and separate more from the DNA in the central tail regions than in H3, where the contacts are more consistent across the tail.

### The intrinsically disordered regions of CENP-A increase overall nucleosomal DNA accessibility

Next, we asked whether we could further refine the region of the histone sequence responsible for DNA conformation. We started by swapping a combination of the two disordered N-terminal regions, the tail and latch, which have the least sequence identity between H3.1 and CENP-A. Surprisingly, we observed that, for both CENP-A_Tail+Latch_H3_Core_ NCP and H3_Tail+Latch_CENP-A_Core_ NCP, DNA is at least as accessible via MNase digestion as the WT CENP-A NCP **(Figure 3A)**. These results suggest that neither the intrinsically disordered N-terminal regions nor the helical core of H3 is sufficient to define the less accessible DNA conformation of the canonical NCP. Interestingly, we observe that the DNA of CENP-A_Tail+Latch_H3_Core_ NCP appears to digest faster than the WT CENP-A NCP.

**Figure 3.**
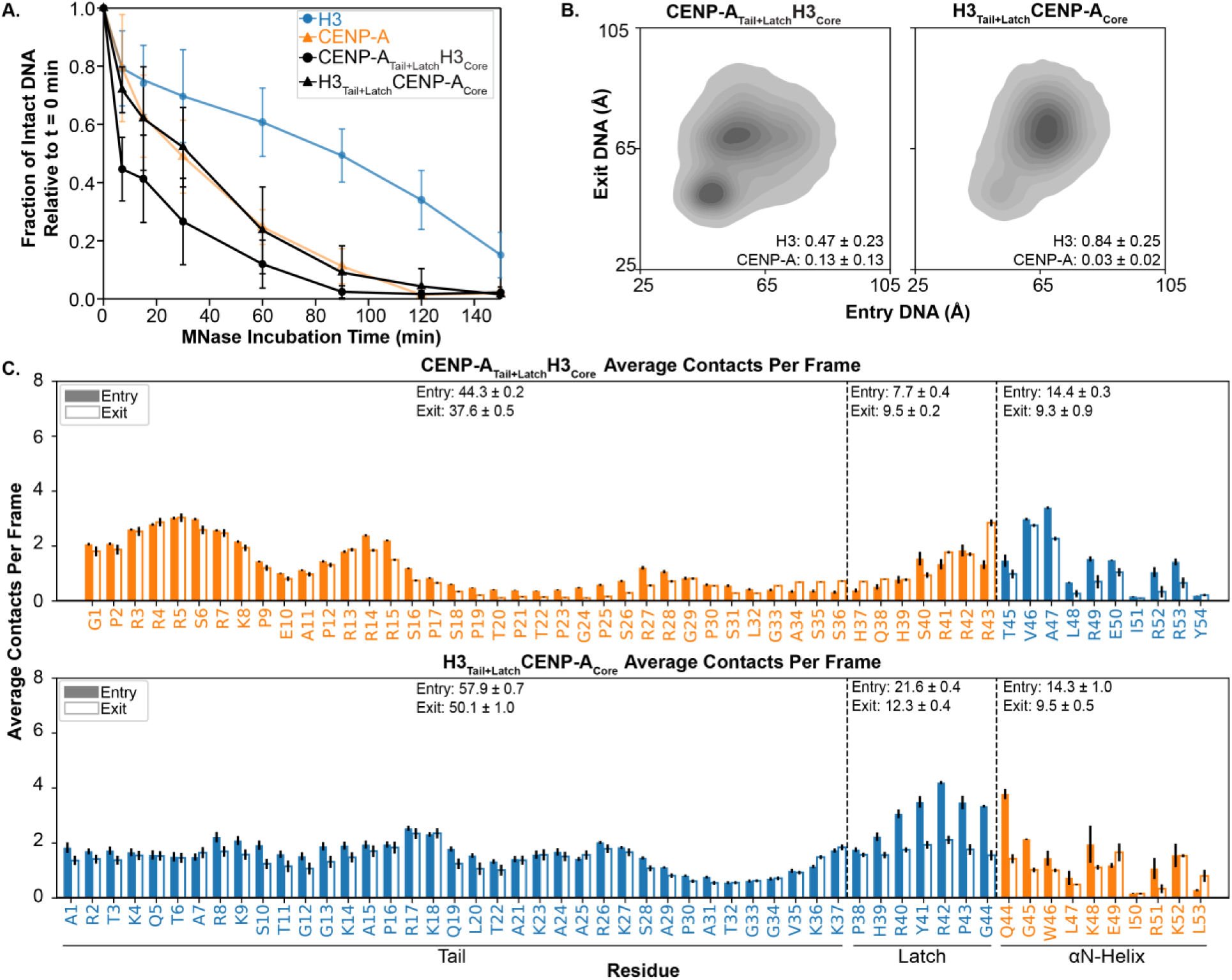
Chimeras swapping the tail and latch fail to swap nucleosomal DNA conformation. **A.** MNase digestion plot of CENP-A_Tail+Latch_H3_Core_ (black circles) and H3_Tail+Latch_CENP-A_Core_ (black triangles) with H3 (blue circles) and CENP-A (orange triangles) for comparison. Plots represent the average of a minimum of n=3 assays, and error bars represent the standard deviation of these data. NCPs use Widom 601 DNA. **B.** Plots of end-to-dyad distances from CG simulations of CENP-A_Tail+Latch_H3_Core_ and H3_Tail+Latch_CENP-A_Core_ NCPs with α-satellite DNA. Plots are inset with KL divergence values, with errors representing standard deviation. **C.** Per-residue bar plots of average contacts with DNA for the tail, latch, and αN-helix regions for CENP-A_Tail+Latch_H3_Core_ and H3_Tail+Latch_CENP-A_Core_ from CG simulations of NCPs with α-satellite DNA. Contacts are shown for entry (solid bars) and exit (open bars) tails. Blue bars and text represent amino acid sequence originating from H3; orange bars and text represent amino acid sequence originating from CENP-A. Per-residue values represent the mean of n=4 independent simulation replicates, with error bars representing the standard error of the mean across replicates. Total average contacts for each region are above the data bars, with errors representing propagated SEM calculated from the individual per-residue SEM values.

Upon examination of the end-to-dyad distances in CG simulations with these two chimeras, we find that the CENP-A tail and latch lead the DNA of CENP-A_Tail+Latch_H3_Core_ NCP to adopt a mixture of open and closed conformations **(Figure 3B)**. The KL divergence values for this NCP relative to H3 and CENP-A were 0.47 ± 0.23 and 0.13 ± 0.13, respectively, demonstrating a greater similarity with a CENP-A-like DNA state. In agreement with the MNase data, the H3 tail and latch are insufficient to reduce DNA accessibility around the CENP-A core in CG simulations of H3_Tail+Latch_CENP-A_Core_ NCP, and the dominant conformation remains open. We note that this chimera yields KL values of 0.84 ± 0.25 relative to H3 and 0.03 ± 0.02 relative to CENP-A, consistent with a DNA conformation similar to that of CENP-A NCP. Thus, H3_Tail+Latch_CENP-A_Core_ NCP primarily samples an accessible conformation while CENP-A_Tail+Latch_H3_Core_ NCP may sample a mixture of open and closed conformations.

Removal of the N-terminal disordered regions in CG simulations provides an interesting point of comparison for these two chimeras. Intriguingly, when the N-terminal disordered regions are removed in CG simulations, the truncated H3 NCP shows increased sampling of open DNA conformations compared to the canonical NCP but is still substantially more closed than with the CENP-A tail and latch (i.e., CENP-A_Tail+Latch_H3_Core_ NCP). In contrast, the H3 tail and latch (i.e., H3_Tail+Latch_CENP-A_Core_ NCP) result in sampling more closed conformations than truncated CENP-A NCP **(Figure S21)**.

Contact profiles for the CENP-A_Tail+Latch_H3_Core_ and H3_Tail+Latch_CENP-A_Core_ NCPs remain broadly consistent with their respective WT origins **(Figures 1E, 3C)**. Tail-DNA contact plots show similar overall profiles to previous systems **(Figures 1E, 2C, 3C)**, while the total H3 tail-DNA contacts continue to decrease with increasing CENP-A composition [entry/exit: 61.8/68.6 (WT) vs. 58.2/57.0 (H3_Tail+Latch+αN_CENP-A_Core_) vs. 57.9/50.1 (H3_Tail+Latch_CENP-A_Core_)]. We also observe a notable successive decrease in the total DNA contacts of the CENP-A latch with increasing H3 composition [entry/exit: 18.1/13.0 (WT) vs. 11.1/11.7 (CENP-A_Tail+Latch+αN_H3_Core_) vs. 7.7/9.5 (CENP-A_Tail+Latch_H3_Core_)]; the total DNA contacts in the CENP-A latch are roughly halved when adjacent to the H3 αN-helix. We speculate that the reduced latch-DNA contacts in CENP-A_Tail+Latch_H3_Core_ NCP are responsible for the high DNA accessibility observed via MNase digestion. Interestingly, the H3 αN-helix exhibits more DNA contacts in CENP-A_Tail+Latch_H3_Core_ NCP than in WT H3 NCP, despite the decrease in latch-DNA interactions, while the CENP-A αN-helix has fewer DNA contacts in H3_Tail+Latch_CENP-A_Core_ NCP than in WT CENP-A NCP.

Overall, we note that the DNA contacts mediated by the tail, latch, and αN-helix are, in part, influenced by the sequence context. Based on the digestive and conformational deviations from the WT NCPs, these chimera swaps demonstrate that the H3 tail and latch are insufficient for promoting a closed NCP conformation but that the CENP-A tail and latch favor an open NCP conformation. Thus, these results delineate a potential cooperative role between the histone tail, latch, and αN-helix in dictating NCP conformation.

### Individual N-terminal regions do not solely dictate overall DNA conformations of canonical and centromeric NCPs

We next sought to determine the minimal region necessary to maintain the DNA conformation of either nucleosome. We created histone chimeras that swapped each region of interest individually, starting with the N-terminal tails. We observed an intermediate MNase digestion profile for CENP-A_Tail_H3_core_ NCP, demonstrating a partial opening of the DNA conformation as compared to WT H3 NCP **(Figure 4A)**. In agreement, our CG simulations of the CENP-A_Tail_H3_Core_ NCP showed noticeable opening of the DNA termini, but they remained firmly wrapped for much of the trajectory **(Figure 4B)**. KL divergence values reveal 0.19 ± 0.10 divergence from H3 NCP and 0.41 ± 0.15 divergence from CENP-A NCP, supporting stronger similarity with H3 NCP. The H3_Tail_CENP-A_Core_ NCP was readily digested by MNase, aligning well with WT CENP-A NCP. In end-to-dyad analysis of H3_Tail_CENP-A_Core_ NCP CG simulations, open DNA termini states are dominant, although a small yet noteworthy population samples more closed DNA termini states. KL values for this NCP are 0.75 ± 0.27 relative to H3 NCP and 0.02 ± 0.08 relative to CENP-A NCP **(Figure 4B, Table S9),** indicating that the DNA conformation is significantly more in line with the CENP-A NCP DNA. When we quantify the total DNA contacts across the N-terminal regions, we observe differences for the CENP-A_Tail_H3_Core_ NCP, including a moderate increase in CENP-A exit tail contacts [entry/exit: 45.6/44.9 vs. 47.2/35.0 (WT)]. Interestingly, when preceded by the CENP-A tail, the H3 latch displays a large decrease in total DNA contacts on the entry side [entry/exit: 7.1/9.0 vs. 20.3/11.7 (WT)], yet the H3 αN-helix-DNA contacts are minimally affected by the substitution of the CENP-A tail [entry/exit: 10.1/7.8 vs. 8.3/9.6 (WT)] **(Figure 4C).** Conversely, in the context of the H3_Tail_CENP-A_Core_ NCP, we see a sharp decline in H3 exit tail contacts as compared to WT H3 NCP [entry/exit: 56.3/43.4 vs. 61.8/68.6 (WT)]. The presence of the H3 tail does not impede CENP-A latch and αN-helix contacts with DNA. Our findings suggest that the CENP-A tail supports the open conformation of the centromeric nucleosome, potentially by decreasing latch-DNA contacts, while the H3 tail sequence is unable to form sufficient DNA contacts to pull the DNA ends closed without its corresponding latch and *a*N sequence.

**Figure 4.**
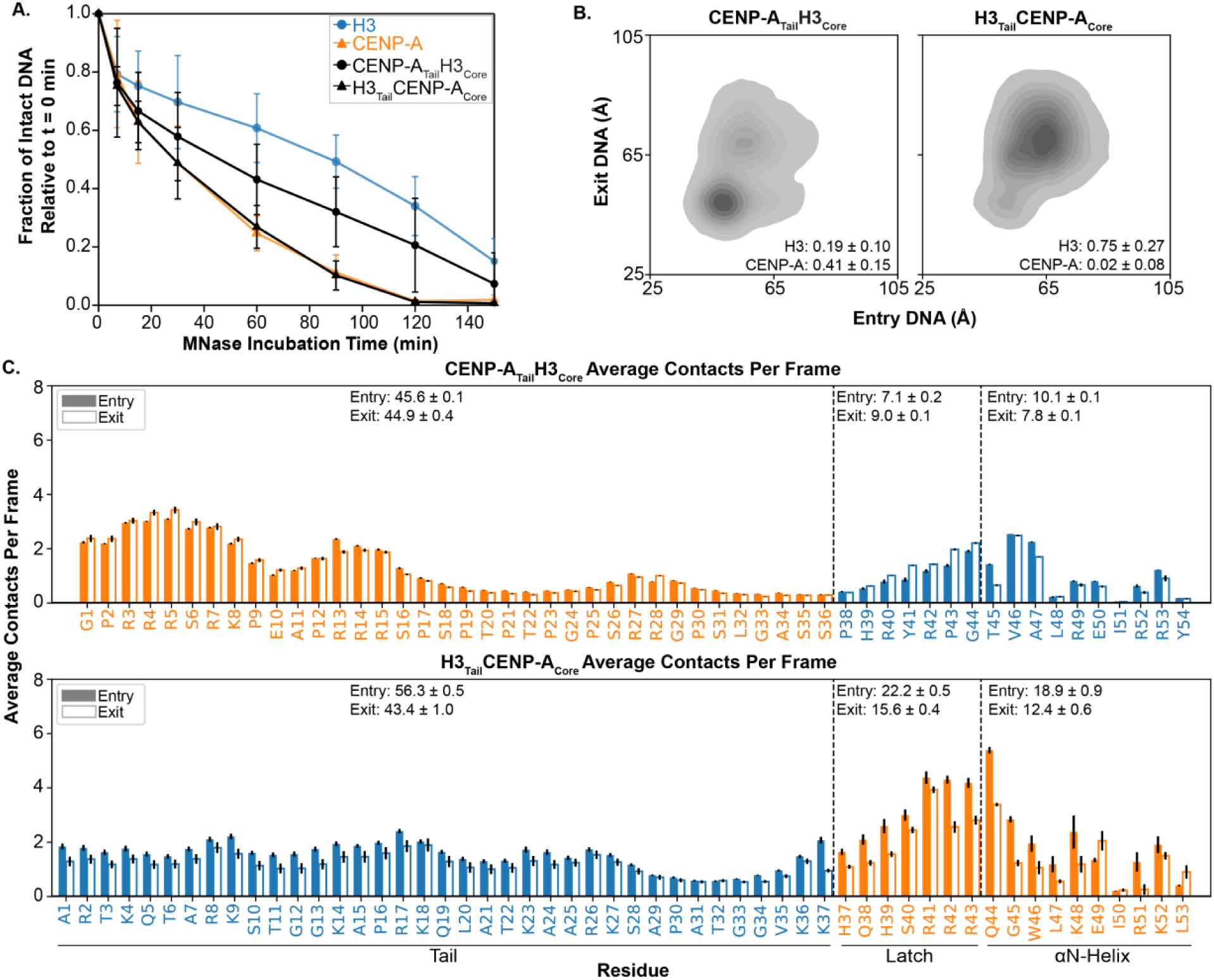
Chimeras swapping the histone tails do not swap nucleosomal DNA conformation. **A.** MNase digestion plot of CENP-A_Tail_H3_Core_ (black circles) and H3_Tail_CENP-A_Core_ (black triangles) with H3 (blue circles) and CENP-A (orange triangles) for comparison. Plots represent the average of a minimum of n=3 assays, and error bars represent the standard deviation of these data. NCPs use Widom 601 DNA. **B.** Plots of end-to-dyad distances from CG simulations of CENP-A_Tail_H3_Core_ and H3_Tail_CENP-A_Core_ NCPs with α-satellite DNA. Plots are inset with KL divergence values, with errors representing standard deviation. **C.** Per-residue bar plots of average contacts with DNA for the tail, latch, and αN-helix regions for CENP-A_Tail_H3_Core_ and H3_Tail_CENP-A_Core_ from CG simulations of NCPs with α-satellite DNA. Contacts are shown for entry (solid bars) and exit (open bars) tails. Blue bars and text represent amino acid sequence originating from H3; orange bars and text represent amino acid sequence originating from CENP-A. Per-residue values represent the mean of n=4 independent simulation replicates, with error bars representing the standard error of the mean across replicates. Total average contacts for each region are above the data bars, with errors representing propagated SEM calculated from the individual per-residue SEM values.

We next investigated the relative contribution of the latch region to nucleosome conformation using additional histone chimeras. Substituting the CENP-A latch into H3 resulted in greater accessibility of the nucleosomal DNA ends to MNase, even to the extent of WT CENP-A NCP. However, the reverse was not true; swapping the H3 latch into CENP-A does not induce closing of the nucleosomal DNA ends **(Figure 5A)**. This suggests an inequivalent role for the latch region in H3 and CENP-A NCPs. The CG simulation end-to-dyad results for these NCPs also demonstrate that the DNA favors open conformations **(Figure 5B)**: KL analyses of both chimeras with respect to WT H3 NCP indicate that they are completely divergent from the canonical nucleosomal DNA (KL values of 1.00 ± 0.02 and 0.99 ± 0.08). Quantification of the contact patterns of the latch-swapped chimera NCPs shows that H3 with the CENP-A latch displays a notable decrease in exit tail-DNA contacts [entry/exit: 58.5/37.2 vs. 61.8/68.6 (WT)] along with a substantial increase in the αN-helix-DNA contacts [entry/exit: 19.1/14.8 vs. 8.3/9.6 (WT)] as compared to WT H3 NCP **(Figure 5C)**. Interestingly, being flanked by the H3 sequence on both sides leads to a moderate increase in latch-DNA contacts of the CENP-A sequence [entry/exit: 19.3/16.6 vs. 18.1/13.0 (WT)]. When we swap the H3 latch into CENP-A, the αN-helix forms fewer DNA contacts than in WT CENP-A NCP [entry/exit: 13.9/8.5 vs. 20.4/12.9 (WT)], and the H3 latch displays a decrease in DNA contacts as compared to WT H3 NCP [entry/exit: 14.5/9.7 vs. 20.3/11.7 (WT)]. Taken together, the CENP-A latch is permissive to an accessible DNA conformation, perhaps by decreasing tail-DNA contacts, but the H3 latch cannot solely dictate a closed conformation.

**Figure 5.**
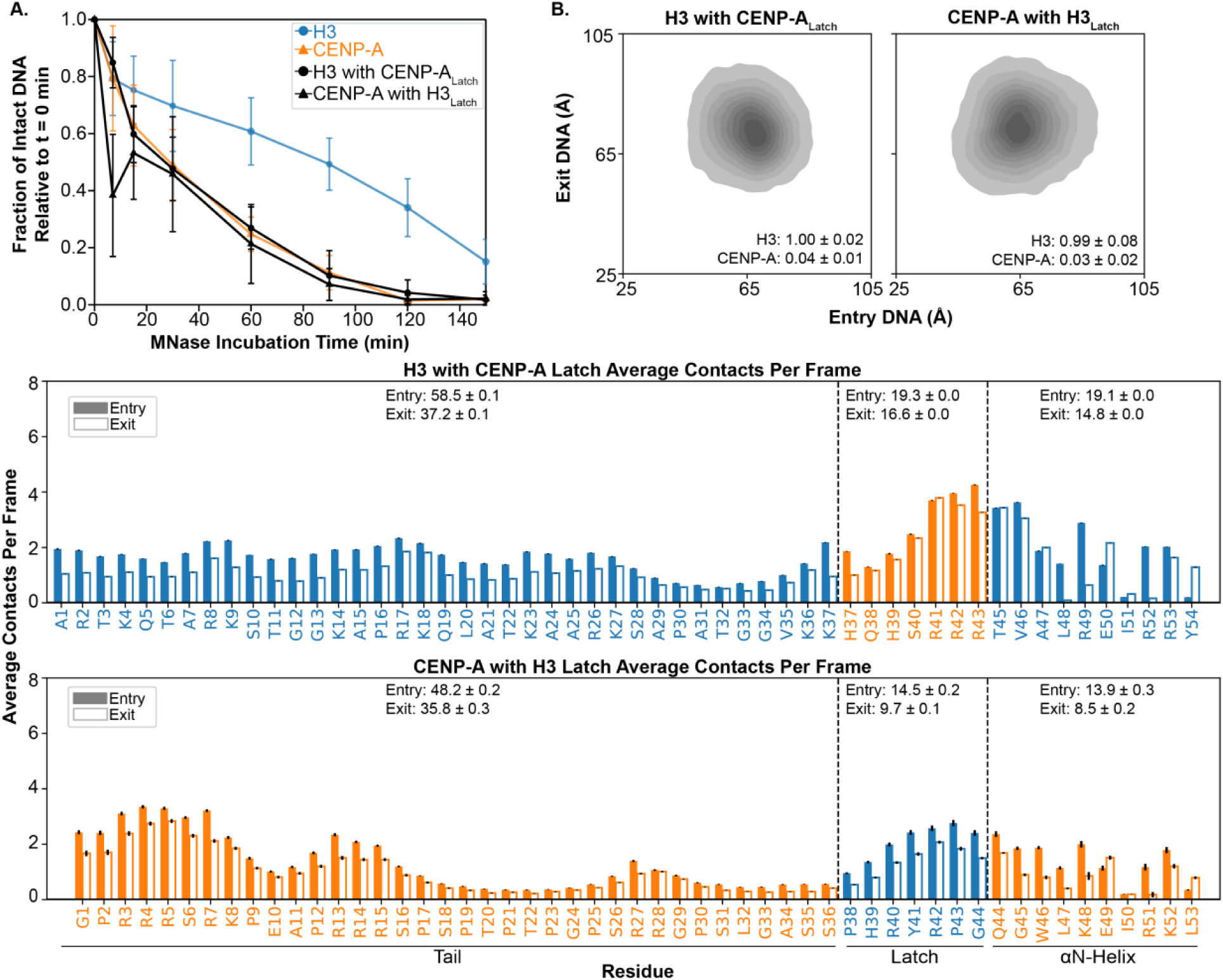
Chimeras swapping the latches support open DNA conformations. **A.** MNase digestion plot of H3 with CENP-A_Latch_ (black circles) and CENP-A with H3_Latch_ (black triangles) with H3 (blue circles) and CENP-A (orange triangles). Plots represent the average of a minimum of n=3 assays, and error bars represent the standard deviation of these data. NCPs use Widom 601 DNA. **B.** Plots of end-to-dyad distances from CG simulations of H3 with CENP-A_Latch_ and CENP-A with H3_Latch_ NCPs with α-satellite DNA. Plots are inset with KL divergence values, with errors representing standard deviation. **C.** Per-residue bar plots of average contacts with DNA for the tail, latch, and αN-helix regions for H3 with CENP-A_Latch_ and CENP-A with H3_Latch_ from CG simulations of NCPs with α-satellite DNA. Contacts are shown for entry (solid bars) and exit (open bars) tails. Blue bars and text represent amino acid sequence originating from H3; orange bars and text represent amino acid sequence originating from CENP-A. Per-residue values represent the mean of n=4 independent simulation replicates, with error bars representing the standard error of the mean across replicates. Total average contacts for each region are above the data bars, with errors representing propagated SEM calculated from the individual per-residue SEM values.

Lastly, we swapped the H3 and CENP-A αN-helix. Our MNase digestion assays show that H3 NCPs with the CENP-A αN-helix have increased DNA accessibility as compared to WT H3 NCP, although not to the extent of WT CENP-A NCP **(Figure 6A)**. Our end-to-dyad calculations from CG simulations agree that the CENP-A αN-helix increases DNA accessibility as compared to WT H3 NCP **(Figure 6B).** However, the KL divergence values from WT H3 and CENP-A NCPs are 0.21 ± 0.11 and 0.30 ± 0.14, respectively, suggesting it is more reminiscent of canonical than centromeric NCP. On the other hand, swapping the H3 αN-helix into CENP-A does not alter the MNase accessibility of the centromeric DNA conformation. In agreement, CG simulations of CENP-A with the H3 αN-helix show similar end-to-dyad distances to WT CENP-A NCP: KL divergence from H3 is 0.86 ± 0.22 for this NCP, while divergence from CENP-A is 0.01 ± 0.02 **(Figure 6B, Table S9)**. In these αN-helix-swapped NCPs, DNA-contact patterns display the greatest change in the αN-helices; the tail and latch regions maintain the characteristic DNA-contact behavior of their respective WT NCPs **(Figure 1E, 5C)**. With one chimera, the substituted CENP-A αN-helix has a reduced contact density relative to WT CENP-A NCP [entry/exit: 7.7/6.0 vs. 20.4/12.9 (WT)] and instead aligns with the lower DNA-contact levels characteristic of the H3 αN-helix in canonical NCP (entry/exit: 8.3/9.6). With the other chimera, the swapped H3 αN-helix shows increased DNA contacts relative to WT H3 NCP [entry/exit: 18.0/16.0 vs. 8.3/9.6 (WT)], similar to the higher contact levels of the CENP-A αN-helix within its WT setting (entry/exit: 20.4/12.9). Interestingly, the number of *a*N-helix-DNA contacts appears to be driven by the sequence N-terminal to the *a*N-helix, which was also observed with the latch-swapped chimeras. Altogether, we observe that neither the H3 αN-helix nor the CENP-A αN-helix can fully recapitulate the respective WT NCP DNA conformations, suggesting the αN-helix is not the sole determinant of NCP conformation.

**Figure 6.**
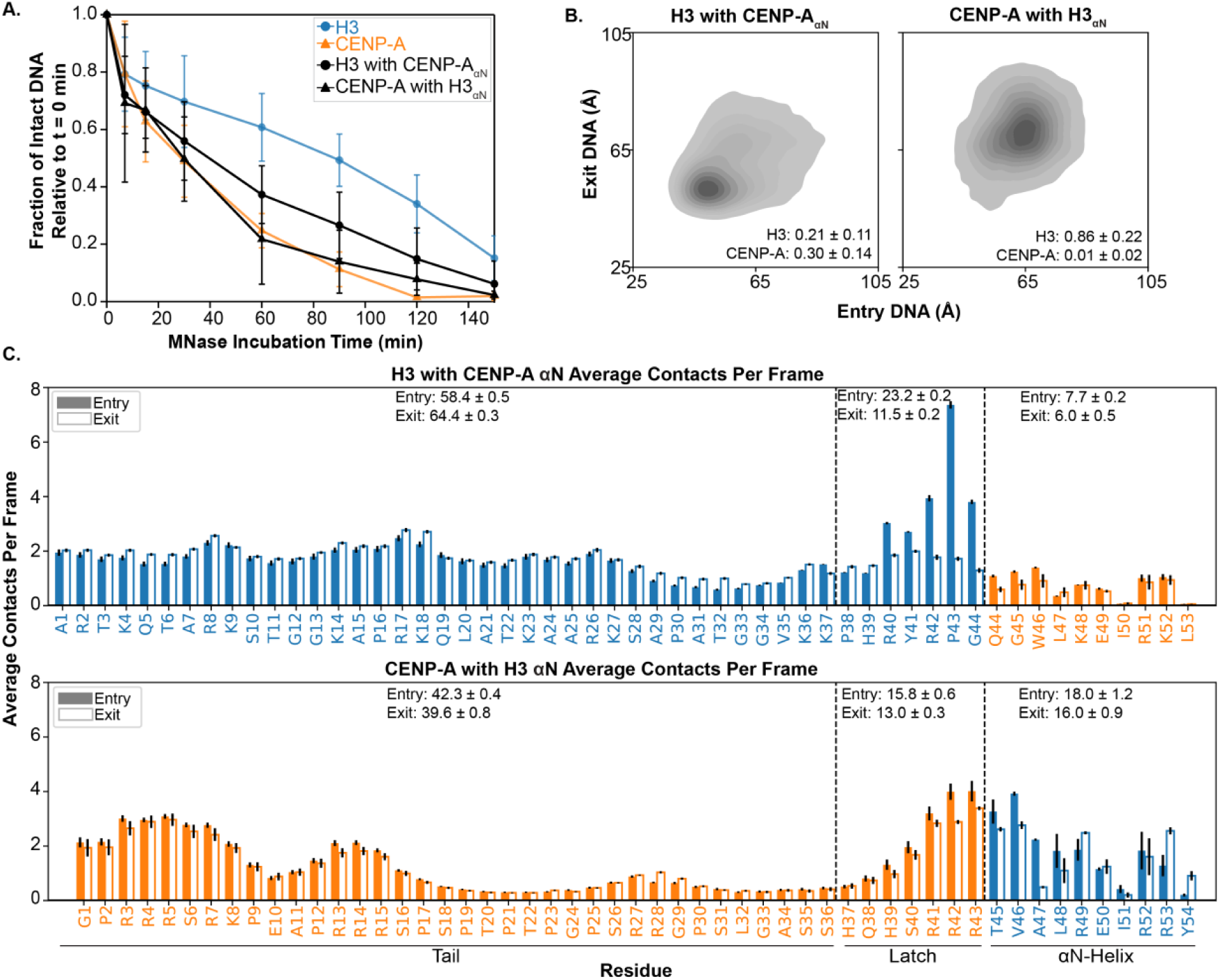
In chimeras, αN swaps are not the sole determinant of nucleosomal DNA conformation. **A.** MNase digestion plot of H3 with CENP-A_αN_ (black circles) and CENP-A with H3_αN_ (black triangles) with H3 (blue circles) and CENP-A (orange triangles). Plots represent the average of a minimum of n=3 assays, and error bars represent the standard deviation of these data. NCPs use Widom 601 DNA. **B.** Plots of end-to-dyad distances from CG simulations of H3 with CENP-A_αN_ and CENP-A with H3_αN_ NCPs with α-satellite DNA. Plots are inset with KL divergence values, with errors representing standard deviation. **C.** Per-residue bar plots of average contacts with DNA for the tail, latch, and αN-helix regions for H3 with CENP-A_αN_ and CENP-A with H3_αN_ from CG simulations of NCPs with α-satellite DNA. Contacts are shown for entry (solid bars) and exit (open bars) tails. Blue bars and text represent amino acid sequence originating from H3; orange bars and text represent amino acid sequence originating from CENP-A. Per-residue values represent the mean of n=4 independent simulation replicates, with error bars representing the standard error of the mean across replicates. Total average contacts for each region are above the data bars, with errors representing propagated SEM calculated from the individual per-residue SEM values.

## DISCUSSION

In this study, we demonstrate that DNA conformations of the canonical and centromeric nucleosomes are not dictated by a single histone region. Through a combination of MNase digestion assays and CG MD simulations conducted on a series of NCPs reconstituted with WT and chimera H3 and CENP-A histones, we show that the H3 tail, latch, and αN-helix are all necessary to adopt the closed DNA conformation of the canonical nucleosome while these three N-terminal regions of CENP-A each contribute to opening the DNA conformation of the centromeric nucleosome. Counterintuitively, we observe a trend across the NCP systems wherein an open DNA conformation correlates with an elevated number of αN-helix-DNA contacts. Conversely, systems with closed DNA conformations have fewer DNA contacts in the αN-helix. This behavior is influenced in part by the identity of the preceding tail and latch region. The presence of the CENP-A latch tends to favor increased contacts in the αN-helix regardless of the sequence context. Histone tail contacts have a less clear role in dictating the overall DNA conformation. However, to maintain a closed conformation like the canonical NCP, the H3 tail, latch, and αN-helix must all be present. Our study demonstrates the regions of the CENP-A and H3 histones that dictate overall nucleosome conformation (**Figure 7**).

**Figure 7.**
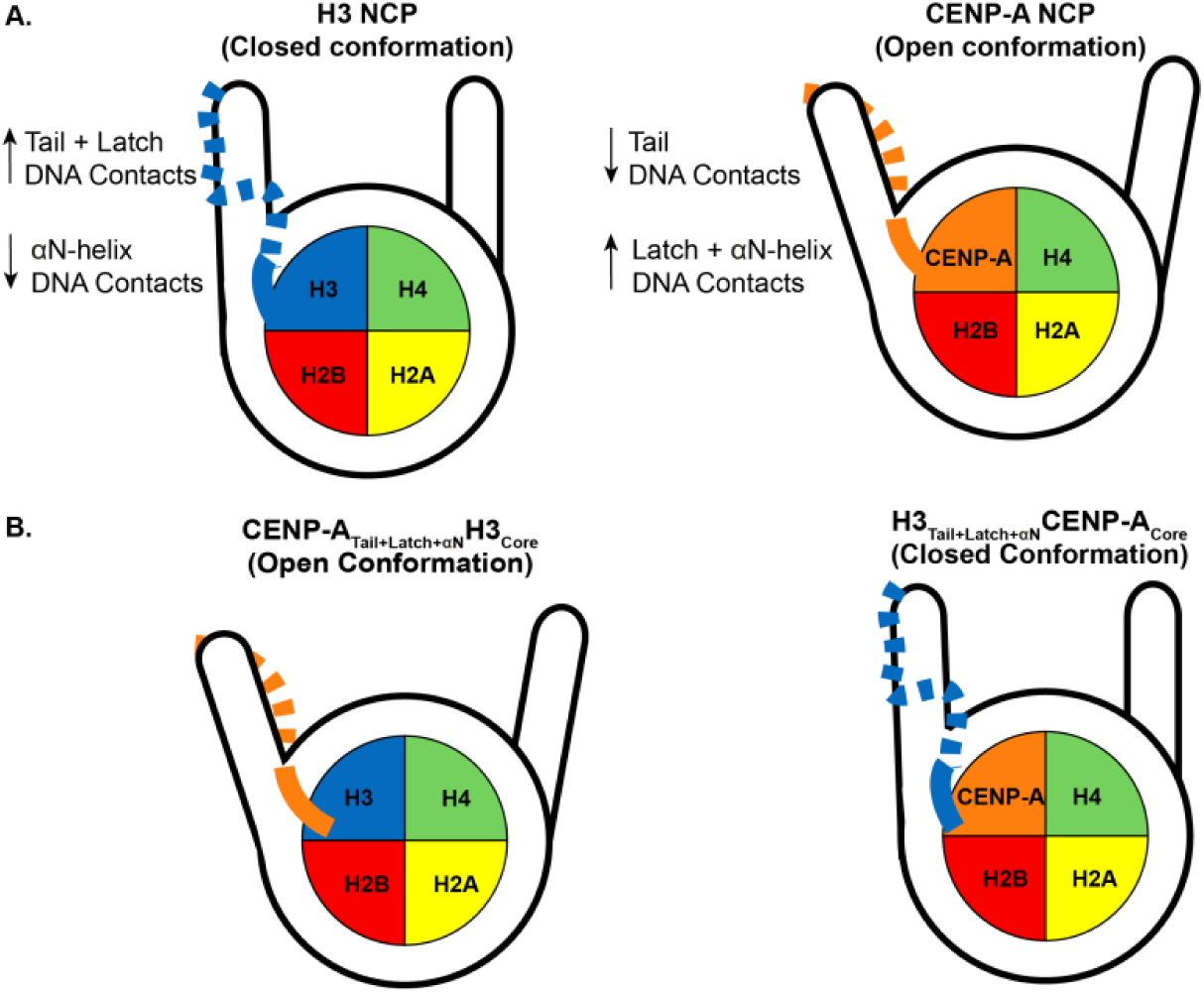
Histone Composition Controls Nucleosome Opening via Coordinated N-terminal region– DNA Interactions. **A.** Schematic representation of nucleosome structural states highlighting the role of histone composition in regulating DNA accessibility. The DNA of the H3 NCP is depicted in a closed conformation, whereas the CENP-A NCP adopts a more flexible, open state. **B.** H3/CENP-A chimeras demonstrate that swapping histone elements modulates this behavior: CENP-A_Tail+Latch+αN_H3_Core_ exhibits a dominant open state, while H3_Tail+Latch+αN_CENP-A_Core_ exhibits a dominant closed configuration. H3- and CENP-A-derived histone domains are color-coded (blue for H3 origin, orange for CENP-A origin) to emphasize the contribution of coordinated interactions between the N-terminal regions and DNA in governing nucleosomal DNA conformation.

Previous simulation studies of CENP-A nucleosomes have lacked extensive sampling or inclusion of the N-terminal tails, limiting conclusions regarding their differential contributions to nucleosome conformation (72–74). This limitation is partly rooted in experimentally determined CENP-A nucleosome structures, which do not resolve the N-terminal tails due to their intrinsically disordered nature (24–27). Consequently, several molecular dynamics studies have either omitted the N-terminal histone-fold extensions entirely or included only the αN-helix before attributing increased DNA end flexibility to the shortened αN-helix of CENP-A (72–74). Our results with full-length histones suggest that the adjacent latch and tail regions play a critical role in modulating DNA conformation and flexibility. Several chimeric constructs exhibit enhanced DNA end dynamics even with the H3 αN-helix, indicating that helix length alone does not fully account for the observed behavior. Rather, we observe that interplay from all three H3 N-terminal components (i.e., the H3_Tail+Latch+αN_CENP-A_Core_ chimera) is required for a closed DNA conformation. Our findings support a more distributed model of regulation, in which local sequence and the context of the adjacent regions within the N-terminal tail-latch-helix collectively contribute to nucleosomal DNA conformation.

Confounding cutoffs of chimeras in the literature caused some confusion over which regions of CENP-A determine the characteristic open DNA conformation. It has been the prevailing paradigm that CENP-A nucleosome conformation is determined by the shortened αN-helix of CENP-A (24, 25). However, the impact of the preceding N-terminal regions has been underappreciated in previous studies. In fact, in one of the key studies supporting the importance of the *a*N-helix, the tested “αN-helix swap” included residues from the “latch” region (25). Although our individual chimera swap of the CENP-A αN-helix into H3 resulted in an NCP with an accessible DNA conformation, inserting the H3 αN-helix into the CENP-A NCP did not create a nucleosome with a closed conformation, suggesting the interactions mediated by the H3 αN-helix alone are insufficient. In addition, a recent study concluded that the CENP-A and H3 NCP DNA conformation is determined by the N-terminal tail; however, the NCP chimera swaps tested to reach this conclusion included residues spanning the N-terminal tail, latch, and first five residues of the αN-helix (26). Indeed, our end-to-dyad analysis and MNase assay of H3_Tail+Latch+αN_CENP-A_Core_ NCP showed that it was the only chimera capable of achieving a majority-closed DNA conformation. The counterpart chimera (i.e., CENP-A_Tail+Latch+αN_H3_Core_) NCP maintained an open DNA conformation as expected. However, our systematic investigation perturbing these regions individually refutes the claim that the conformation is dictated by the histone tail. We observe that our CENP-A_Tail_H3_Core_ NCP has an intermediate conformation with a partial opening of the DNA conformation compared to WT H3 NCP, but the H3_Tail_CENP-A_Core_ NCP is as accessible as WT CENP-A NCP. Therefore, while the histone tail is not the sole key determinant, we suggest that the differences in DNA contacts mediated by H3 and CENP-A histone entry/exit tails work in tandem with the other N-terminal regions to maintain overall conformation, as previously suggested (75).

Notably, incorporation of any N-terminal CENP-A histone region increases the accessibility of the NCP. We speculate that this intrinsic property of the CENP-A histone sequence ensures that the centromeric nucleosome maintains the necessary conformational ensemble for CCAN assembly and function in mitotic chromatin (26, 36). Similarly, this redundancy could ensure that the overall CENP-A DNA conformation remains flexible regardless of PTMs such as tri-methylation (76) and phosphorylation (77) of the N-terminal tail, as well as other regulatory modifications in the core (74, 78). On the other hand, the H3 histone is heavily modified across the entire histone (79). Perhaps the combination of the H3 tail, latch, and αN-helix strongly promotes a closed conformation to allow these PTMs to tune nucleosomal DNA accessibility. Ultimately, consideration of the intrinsically disordered histone regions is necessary to fully understand nucleosome DNA conformation. The unique structural and dynamic properties of the CENP-A nucleosome may reconcile the dichotomy between the observed local flexibility of the CENP-A nucleosome and the chromatin compaction promoted by CENP-A.

## DATA AVAILABILITY

The data underlying this article are available in the manuscript, the supplementary materials, and upon reasonable request to the appropriate corresponding author. Simulation analyses scripts are available on GitHub (https://github.com/WereszczynskiGroup/supplemental-data-cenpa-2026). Raw trajectory data generated for this study are available on Zenodo at (https://doi.org/10.5281/zenodo.22307430). All-atom trajectories are stripped of solvation and temporally strided to reduce file size but still cover the full duration of simulation. Together, with instructions found on the GitHub, these resources are sufficient to reproduce all analyses and results reported in this manuscript and the corresponding SI.

## ACKNOWLEDGEMENTS

This work was supported by the National Institutes of Health: grant R35GM142594 to E.A.M and R35GM119647 to J.W. Supported, in part, by Institutional Research Grant IRG #19-138-34 from the American Cancer Society and the MCW Cancer Center. This manuscript is subject to the NIH Public Access Policy. Through acceptance of the federal funding used to support the research reported in this manuscript, the NIH has been given a right to make this manuscript publicly available in PubMed Central upon the Official Date of Publication, as defined by the NIH. The funders had no role in study design, data collection and analysis, decision to publish, or preparation of the manuscript. The content is solely the responsibility of the authors and does not necessarily represent the official views of the National Institutes of Health. Thank you to Dr. Aaron Straight for the bicistronic CENP-A/H4 plasmid. Thank you to Drs. Catherine Musselman, Michael Poirier, and Karolin Luger for histone plasmids. Thank you to the MCW Department of Biochemistry for access to the mass spectrometer.

## AUTHOR CONTRIBUTIONS

EM developed the initial concept of the project. EM and JW obtained funding and supervised the project. EP and SMZ prepared samples. EP and JL conducted the research, analyzed, and visualized the data. EP and JL wrote the manuscript, with contributions and edits from EM and JW.

## SUPPORTING MATERIAL

Supporting material is available online.

