## Supporting Information for "Determinants of centromeric nucleosome conformation"

### Supplemental Methods

***All-atom nucleosome construction.*** The canonical nucleosome structures of H3 (H3.1 C96S/C110A) and CENP-A were initially obtained from Robetta using the 1KX5 crystal structure of the canonical eukaryotic nucleosome as a template with the Widom 601 DNA sequence to build similar initial structures in closed DNA conformations (1–3). Chimeric systems were built using the UCSF Chimera interface to swap the tails and latches at provided residue cutoffs of $H3^{1\text{-}44}$/CENP-$A^{44\text{-}139}$ and CENP-$A^{1\text{-}43}$/$H3^{45\text{-}135}$ (4). Chimera was also used to delete the tails for tailless nucleosomes to obtain $H3^{45\text{-}135}$ and $CENPA^{44\text{-}139}$ systems. The H3 and CENP-A N-terminal tails of all tailed systems were manually extended away from the nucleosome cores using Chimera to remove initial tail binding bias from Robetta. A second set of all-atom systems was made using a centromeric α-satellite DNA (**see Main Text Materials and Methods**).

***All-atom MD simulation setup and parameters.*** All atom systems were solvated and neutralized with 0.150 M KCl via the tleap and ParmED packages in AmberTools (5–7). The Amber ff19SB (8) , BSC1 (9) , OPC (10) , and Li and Merz (11, 12) 12-6 OPC water ions force fields were used to parameterize the protein, DNA, water, and ions, respectively. Hydrogen mass repartitioning was used to allow for a 4-fs timestep (13). All systems were energy-minimized for 10,000 steps twice, switching from the steepest descent method to the conjugate gradient method both times after 5,000 steps. The first minimization applied 10 kcal/mol/$Å^2$ harmonic restraints to all heavy atoms. The second minimization did not have any restraints. Following minimization, systems were gradually heated from 5 K to 300 K using a Berendsen barostat (14) under Langevin dynamics (15) in an NPT ensemble over the course of 100 ps using the same heavy atom restraints. Heavy atom restraints were slowly lifted in factors of 1/3 every few hundred ps with the first three relaxations lasting 100 ps, the next three rounds 200 ps, and the final round lasting 1000 ps in an NPT ensemble. Complete removal of restraints occurred at 900 ps out of the total 1900 ps. Four replicas per system were run for production simulations for 2.3 μs each using the GPU-accelerated pmemd.cuda in AMBER (7, 16–18). All production runs were in the NPT ensemble, at 300 K, and used a 10 Å non-bonded cutoff along with Particle Mesh Ewald treatment for longer ranged electrostatics (19). The first 300 ns of production time for each replica was dedicated as 'equilibration' time to allow tails to fall onto the nucleosome. Solvation box sizes were adjusted after equilibration time to avoid periodic interactions caused by system size increases **(Figure S7)**. After the addition of a new water box, another round of minimization, heating, and relaxation followed exactly as previously described to adjust the new water box. The remaining 2 μs of production time was used for post-trajectory analysis, for a total of 8 μs post-equilibration time per system **(Table S1)**. Production trajectories were saved every 10 ps and visualized using Visual Molecular Dynamics (VMD) (20). A mix of Amber 20 and Amber 22 was used to run minimization, heating, relaxation, and production for Widom DNA systems as we shifted using versions (7). Amber 22 was used exclusively for centromeric systems.

***All-atom-specific MD simulation energetics analysis.*** The molecular mechanics generalized born surface area (MM/GBSA) energetic calculation analyses of all-atom simulations were performed using the MMPBSA.py program (21, 22) with igb (23, 24) = 8 and a defined salt concentration of 0.150 M. A single-trajectory approach was used where the DNA was defined as the receptor, specific protein residue selections defined as the ligand. All molecular dynamics simulations and conformational sampling were performed in the context of the complete NCP, and components outside of the DNA and protein selections of interest were stripped only during post-processing of the resulting trajectories for MM/GBSA analysis. The remaining DNA and protein selection constituted the complex for each calculation. The same residue selections used to evaluate contacts of the tail and αN-helices with the DNA were used for all relevant MM/GBSA calculations. Residue decomposition calculations within MMPBSA.py were used to obtain the energetic contributions of individual protein residues for residue-based MM/GBSA analyses. In-house python scripts were used to calculate errors for all MM/GBSA analyses and obtain residue decomposition visuals. The python timeseries tool for the multistate Bennett acceptance ratio (PyMBAR) was used to compute the decorrelation time for each individual complex calculation (25, 26).

### Supplemental Results

#### Comparison of all-atom and coarse-grained simulations.

We conducted all-atom MD simulations of the canonical H3 and CENP-A nucleosomes to gain atomic-level insight into the specific intermolecular interactions that dictate nucleosomal conformation. One set of all-atom models was constructed with Widom 601 DNA **(Figures S8, S10-S11, S14A, S15A),** and another with a centromeric α-satellite DNA sequence **(Figures S9, S12-S13, S14B, S15B)**. Each all-atom system was simulated four times for 2 μs per replica post-equilibration, for a total of 48 μs of simulation time for each DNA sequence (96 μs overall). The N-terminal tails of all tailed nucleosome systems collapsed onto the DNA during the 300 ns dedicated equilibration time in each replica.

To assess the favorability of histone-DNA interactions, we conducted MM/GBSA analyses on this all-atom dataset to calculate binding energetics of the N-terminal tail, latch, and αN-helix regions of the H3 and CENP-A histones **(Figure S8-S9, Table S3-S8)**. The MM/GBSA calculations presented here are intended to provide qualitative estimates of relative interaction energetics and should not be interpreted as rigorous, absolute binding free energies. In particular, limitations including incomplete treatment of configurational entropy, solvent approximations, and sensitivity to conformational sampling may affect the quantitative accuracy of the calculated energies. We observe that across this entire region, CENP-A and H3 histones exhibit similar DNA binding energies with Widom 601 DNA, with CENP-A showing slightly more favorable overall interactions **(Figure S8)**. In the disordered N-terminal region, the WT CENP-A tails have more favorable interaction energies relative to the WT H3 tails with Widom DNA (entry/exit: -78.2/-89.8 vs. -57.4/-65.2 kcal/mol) In the latch region, the H3 has slightly more energetically favorable interactions in the WT NCPs relative to CENP-A (entry/exit: -38.3/-40.6 vs. -37.2/-33.6 kcal/mol), and this continues through the αN-helix (entry/exit: -22.3/-18.0 vs. -8.6/-9.6 kcal/mol). The same observational trends are evident from Widom 601 all-atom hydrogen bonds and contacts analyses **(Figure S10A,C)**. Taken together, this suggests there are thermodynamic differences between H3 and CENP-A interactions with DNA across the N-terminal regions.

To further explore the differences between H3 and CENP-A NCPs, we analyzed end-to-dyad distances of both the entry and exit DNA termini from the dyad bp. In the Widom 601 all-atom simulations, CENP-A exhibits a very compact, closed DNA conformation akin to the H3 NCP **(Figure S10B)**. These more compact conformations in CENP-A all-atom simulations were inconsistent with the increased accessibility observed in our MNase experiments **(Figure 1B)** and other previously determined CENP-A structures.

This discrepancy in compaction led us to hypothesize that the shorter timescales inherent to all-atom simulations were preventing the CENP-A DNA from opening (27). We therefore transitioned to conducting coarse-grained (CG) simulations of the canonical H3 and centromeric NCPs with α-satellite DNA to test whether we could observe flexible DNA opening in the CENP-A nucleosome on longer timescales (see **Main Text Results**). The adjustment from all-atom to CG simulations demonstrated that the longer timescales accessible to CG methods are necessary to sample the open conformations of the CENP-A NCP. In an effort to compare the all-atom simulations to the equivalent timeframe of the CG simulations, we performed a time-based end-to-dyad analysis for the initial steps of the CG simulations that were calculated to be the estimated equivalent of the 2 μs of the all-atom simulations. This comparison showed that early CG simulations behave similarly to the all-atom simulations **(Figure S16)**. We therefore used CG models for the remainder of our study.

Importantly, the differences between all-atom and CG simulations reflect distinct modes of tail-DNA engagement rather than contradictory trends in interaction. In all-atom simulations, histone tails predominantly remain positioned along the top, exterior surface of the nucleosome DNA **(Figure S17)**. In contrast, CG simulations allow increased penetration of the tails into the nucleosome interior, between the DNA arms, leading to more frequent inner contacts facing the histone core and altered contact distributions **(Figure S18)**. These differences in spatial sampling provide a structural explanation for the observed variations in contact magnitudes between all-atom and CG simulations, particularly for the H3 and CENP-A NCPs.

#### In CG simulations, the CENP-A αN-helix is associated with greater flexibility in the latch and tail regions.

To examine whether differences in DNA accessibility correlate with changes in histone flexibility, we calculated residue-level root mean square fluctuations (RMSF) for the N-terminal regions of H3 and CENP-A across all CG simulations. Both H3 and CENP-A NCPs exhibit comparable levels of flexibility, with RMSF values ranging from approximately 12-20 Å in the tails that gradually decrease through the latch and into the αN-helix. This overall pattern is consistent across all nucleosomes, indicating that the intrinsic mobility of the histone tails is largely conserved.

While tail fluctuations are similar across NCPs, differences are observed within the latch and αN-helix regions. NCPs containing the CENP-A αN-helix display elevated RMSF values in the adjacent latch region compared to those containing the H3 αN-helix. For example, the CENP-A and CENP-$A_{Tail+Latch+\alpha N}H3_{Core}$ NCPs exhibit increased fluctuations across the latch-αN-helix regions relative to the canonical H3 NCP **(Figure S19)**. In contrast, NCPs containing the H3 αN-helix, including the canonical H3 NCP, show reduced RMSF values across this region.

Chimeric systems with individual component swaps display intermediate RMSF profiles that follow the identity of the latch and αN-helix. Replacement of the H3 αN-helix with the CENP-A αN-helix in an H3 core NCP increases RMSF in both the helix and adjacent latch regions compared to the canonical H3 system. Similarly, substitution of the CENP-A latch into an H3 core NCP increases fluctuations within the latch region. Conversely, the $H3_{Latch+\alpha N}$CENP-$A_{Core}$ NCP shows reduced RMSF values across this latch-helix region compared to the CENP-A NCP **(Figure S19)**.

Tail-only swaps produce comparatively small changes in RMSF profiles outside of the tail region. The CENP-$A_{Tail}H3_{Core}$ NCP and $H3_{Tail}$CENP-$A_{Core}$ NCPs both display tail RMSF profiles similar in magnitude and shape to their respective unperturbed NCPs, while differences remain localized to the latch-αN-helix regions.

Across all systems, RMSF values are highest in the N-terminal tails and decrease through the latch into the αN-helix. Variations in flexibility between NCPs are most pronounced in the latch-αN-helix region, whereas tail flexibility remains largely consistent.

**Distinct Tail–DNA Interaction Patterns Underlie Divergent DNA Conformations.**

We asked if the consistent contacts of the H3 tail residues along the DNA are a driving factor in the closed conformation of the H3 NCPs. To further investigate the impact of histone tail-DNA interactions on varying DNA accessibility across our chimera NCPs, we measured cumulative contacts between the H3/CENP-A N-terminal tails and DNA across all 147 bp to assess the dispersion of contacts along the nucleosomal DNA. We observed appreciable differences between the WT H3 and CENP-A NCPs. Although the H3 NCP has more tail-DNA contacts overall, the CENP-A tail exhibits greater dispersion, contacting a broader range of DNA positions. Across all NCPs, cumulative contact profiles remain highly linear ($R^2$ roughly 0.93-0.97), indicating that contact accumulation is structured and consistent along the DNA length **(Figure S20)**.

To better resolve how these contacts are distributed, we separated them based on DNA orientation relative to the histone core, distinguishing inward-facing from outward-facing DNA. Inside contacts exhibit slightly higher linearity ($R^2$ roughly 0.95-0.97), consistent with a conserved and constrained contact network, while outside contacts show modestly lower linearity ($R^2$ roughly 0.91-0.94), indicating increased variability. While NCPs with increased DNA accessibility generally contain CENP-A cores that exhibit reduced cumulative contacts, contact magnitude alone does not define DNA conformation. NCPs can retain high inside DNA contacts while still sampling open DNA conformations.

Importantly, these results reconcile with earlier per-residue contact analyses, which showed that the H3 tail residues form broadly distributed contacts along the tail length, whereas CENP-A contacts appear localized to arginine-rich regions. While H3 displays greater dispersion along the tail sequence, these interactions are focused on more restricted regions of the DNA. In contrast, the CENP-A tail, despite having more localized contacts along the tail length, engages a wider range of DNA positions. Together, these findings suggest that DNA accessibility is not only governed by the total number of contacts being made, but by their spatial organization along the DNA, which is further reflected in differences in tail positioning observed between all-atom and CG simulations. H3 tails favor persistent, localized interactions, and CENP-A promotes more distributed and flexible engagements.

### Supplemental Figures.

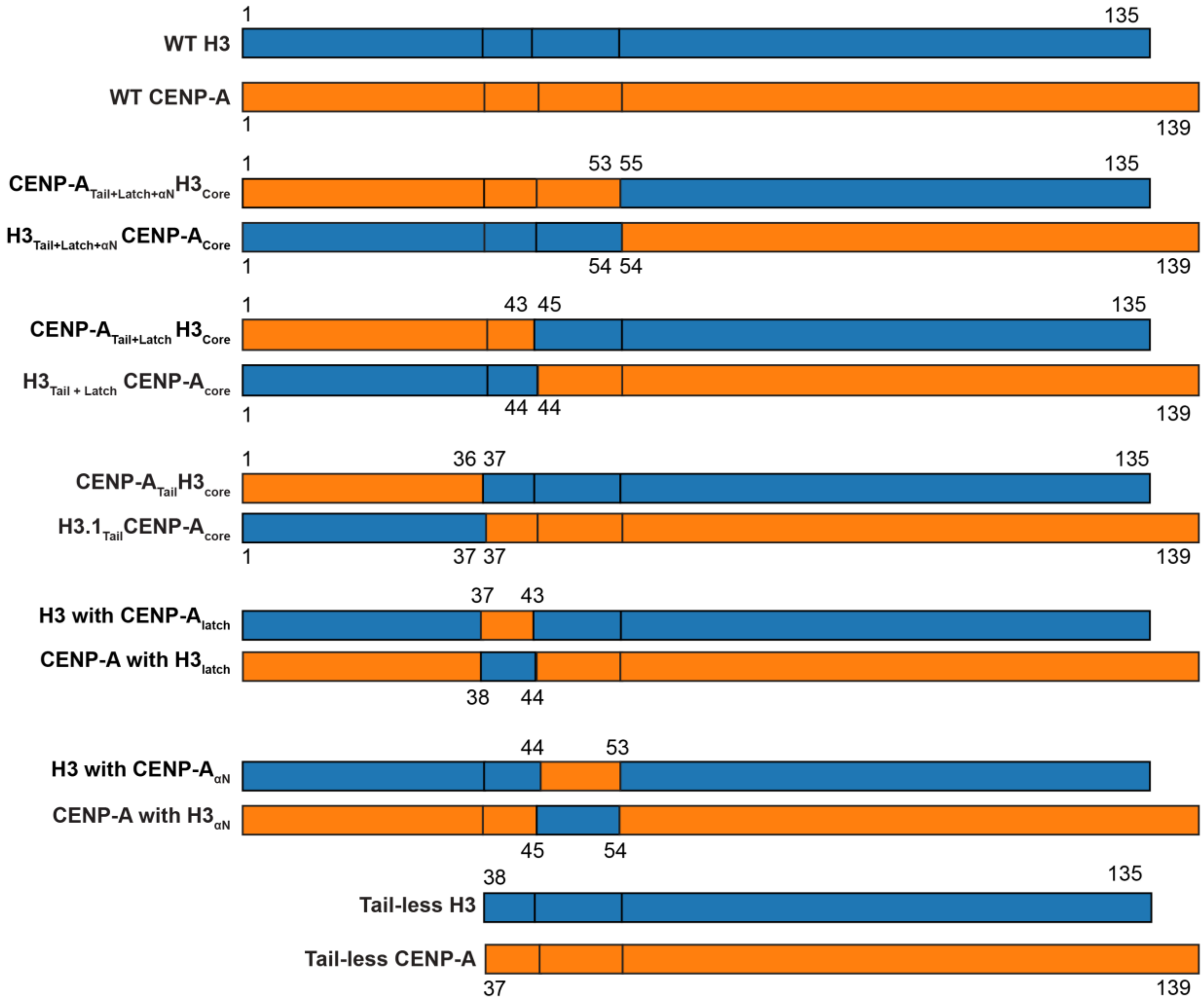


**Figure S1. Chimera Histone Constructs.** Schematic of H3-CENP-A chimera histones reconstituted into NCPs in the study. Blue represents regions from H3 and orange from CENP-A.

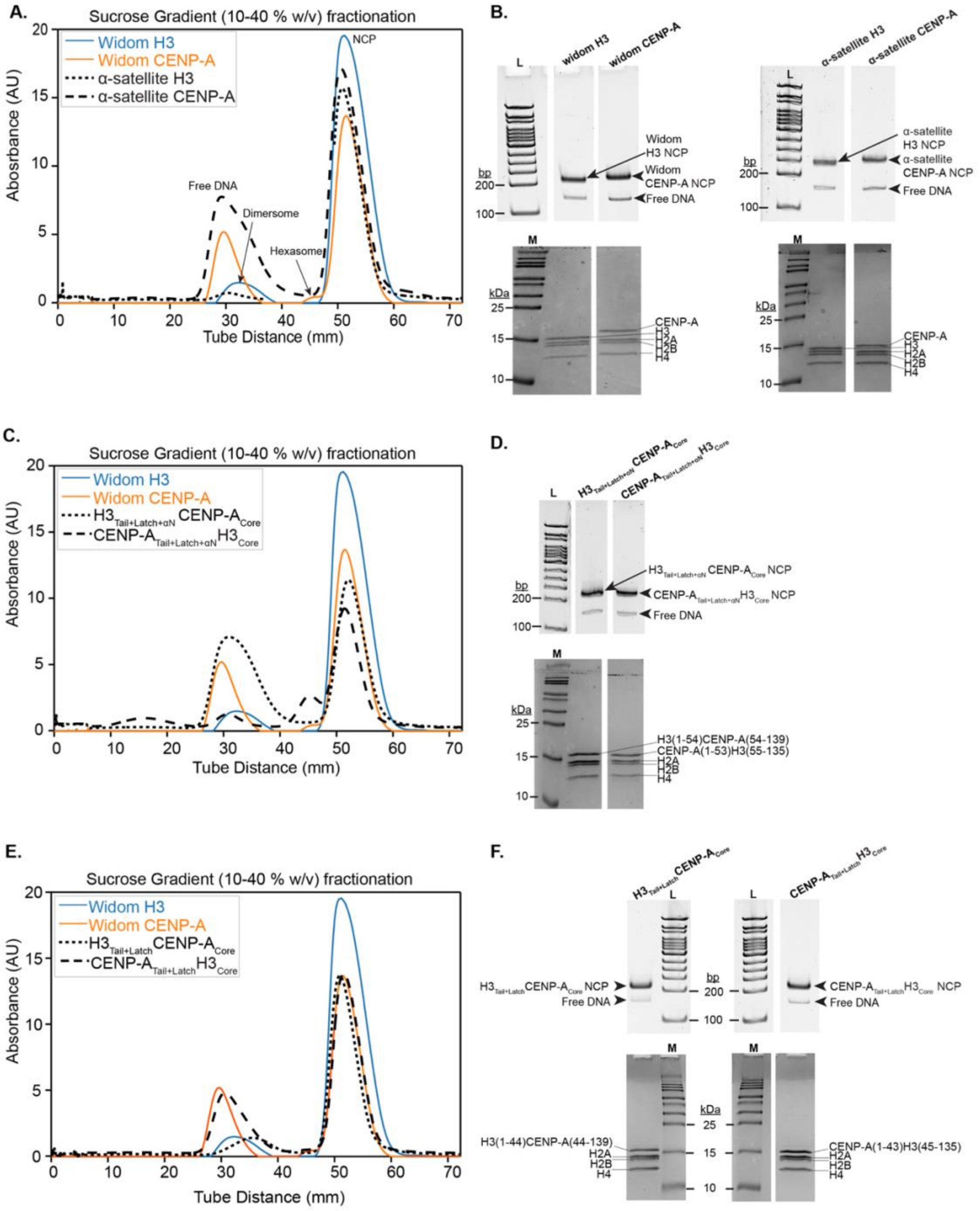


**Figure S2. Validation of WT and Chimera NCPs. A.** Sucrose gradient trace overlay with Widom H3 (blue) Widom CENP-A (orange), α-satellite H3 (black dotted line) and α-satellite CENP-A (dashed line). **B.** 5% Native-PAGE (top) and 18% SDS-PAGE (bottom) of the Widom H3 and CENP-A, and α-satellite H3 and CENP-A of H3 and CENP-A NCPs. In all gels “L” denotes the TrackIt 100 bp DNA ladder (ThermoFisher) and “M” denotes the SpectraBR Protein marker (ThermoFisher). **C.** Sucrose gradient trace overlay with Widom H3 (blue) Widom CENP-A (orange), $H3_{Tail+Latch+\alpha N}$CENP-$A_{Core}$ NCP (black dotted line) and CENP-$A_{Tail+Latch+\alpha N}H3_{Core}$ NCP (dashed line) **D. 5**% Native-PAGE (top) and 18% SDS-PAGE (bottom) of $H3_{Tail+Latch+\alpha N}$CENP-$A_{Core}$ and CENP-$A_{Tail+Latch+\alpha N}H3_{Core}$ NCPs**. E.** Sucrose gradient trace overlay with Widom H3 (blue) Widom CENP-A (orange), $H3_{Tail+Latch}$CENP-$A_{Core}$ (black dotted line) and CENP-$A_{Tail+Latch}H3_{Core}$ (dashed line) **F.** 5% Native-PAGE (top) and 18% SDS-PAGE (bottom) of $H3_{Tail+Latch}$CENP-$A_{Core}$ and CENP-$A_{Tail+Latch}H3_{Core}$ NCPs.

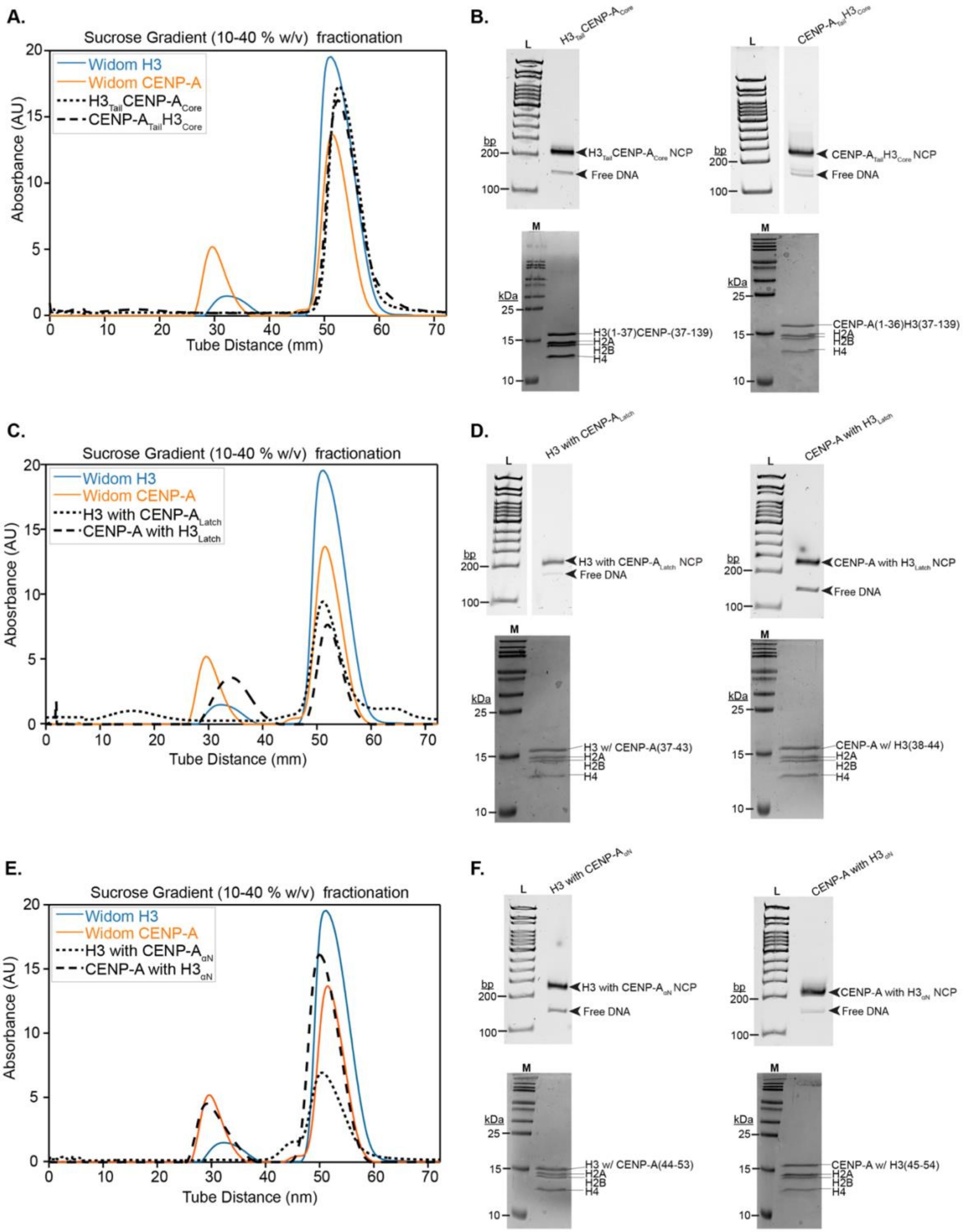


**Figure S3. Validation of Individual Swap Chimera NCPs. A.** Sucrose gradient trace overlay with Widom H3 (blue) Widom CENP-A (orange), H3$_{Tail}$CENP-A$_{Core}$ NCP (black dotted line) CENP-A$_{Tail}$H3$_{Core}$ NCP (dashed line). **B.** 5% Native-PAGE (top) and 18% SDS-PAGE (bottom) of the H3$_{Tail}$CENP-A$_{Core}$ and CENP-A$_{Tail}$H3$_{Core}$ NCPs. . In all gels "L" denotes the TrackIt 100 bp DNA ladder (ThermoFisher) and "M" denotes the SpectraBR Protein marker (ThermoFisher). **C.** Sucrose gradient trace overlay with Widom H3 (blue) Widom CENP-A (orange), H3 with CENP-A$_{Latch}$ (black dotted line) and CENP-A with H3$_{Latch}$ (dashed line). **D.** 5% Native-PAGE (top) and 18% SDS-PAGE (bottom) of the H3 with CENP-A$_{Latch}$ and CENP-A with H3$_{Latch}$ NCPs. **E.** Sucrose gradient trace overlay with Widom H3 (blue) Widom CENP-A (orange), H3 with CENP-A$_{\alpha N}$ (black dotted line) and CENP-A with H3$_{\alpha N}$ (dashed line). **F.** 5% Native-PAGE (top) and 18% SDS-PAGE (bottom) of the H3 with CENP-A$_{\alpha N}$ and CENP-A with H3$_{\alpha N}$ NCPs.

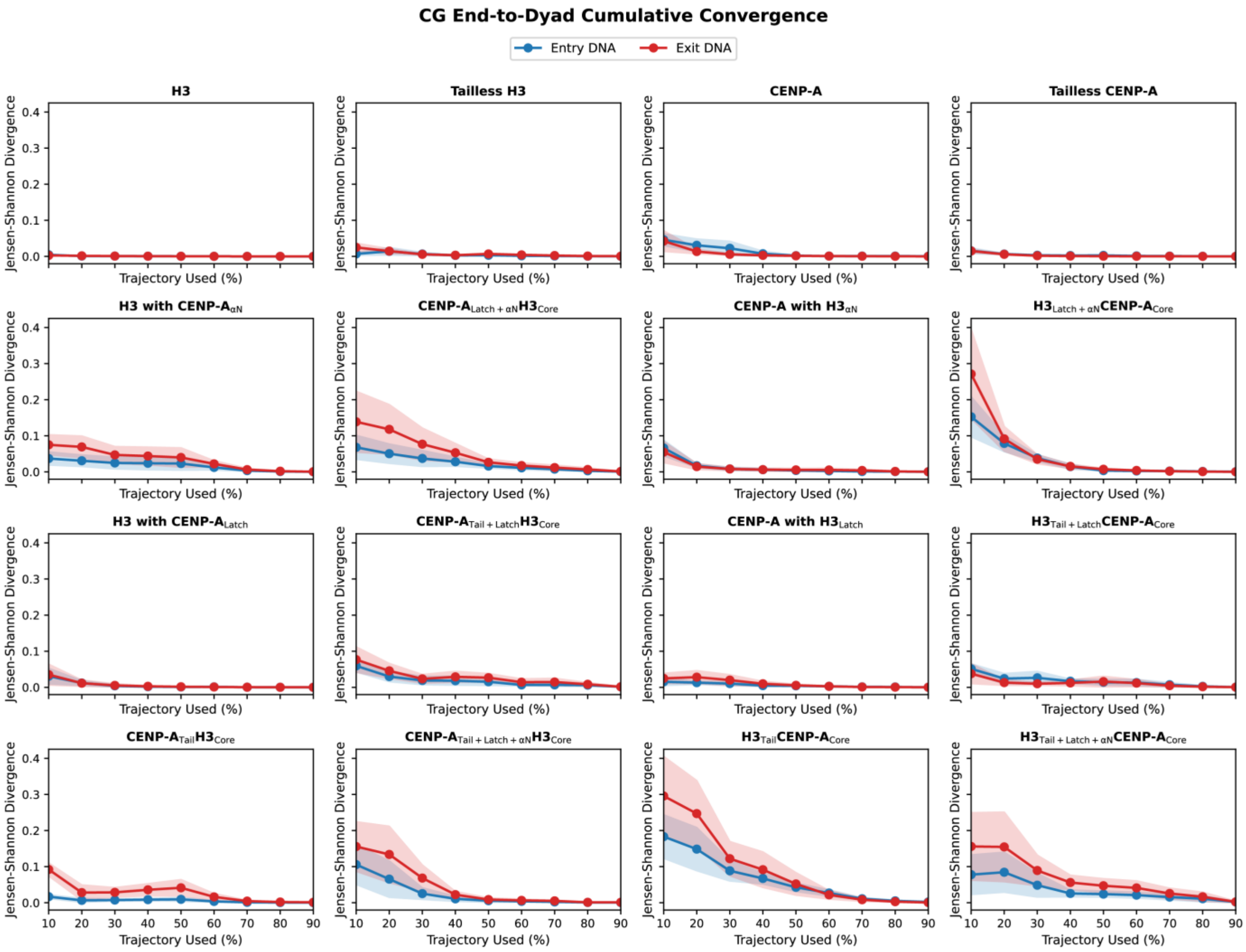


**Figure S4. Convergence Analysis of Coarse-Grained End-to-Dyad Distributions.** Convergence of coarse-grained alpha-satellite DNA nucleosome opening was assessed by calculating EtD distributions using progressively larger fractions (10-90%) of each production trajectory and comparing each cumulative distribution with the corresponding full-trajectory distribution using Jensen-Shannon divergence (JSD). Entry and exit DNA are shown separately. Lines represent the mean JSD across four independent simulation copies, with shaded regions representing the standard error of the mean (SEM, n=4). Lower JSD values indicate greater similarity to the full-trajectory EtD distribution and stabilization of the sampled nucleosome-opening behavior.

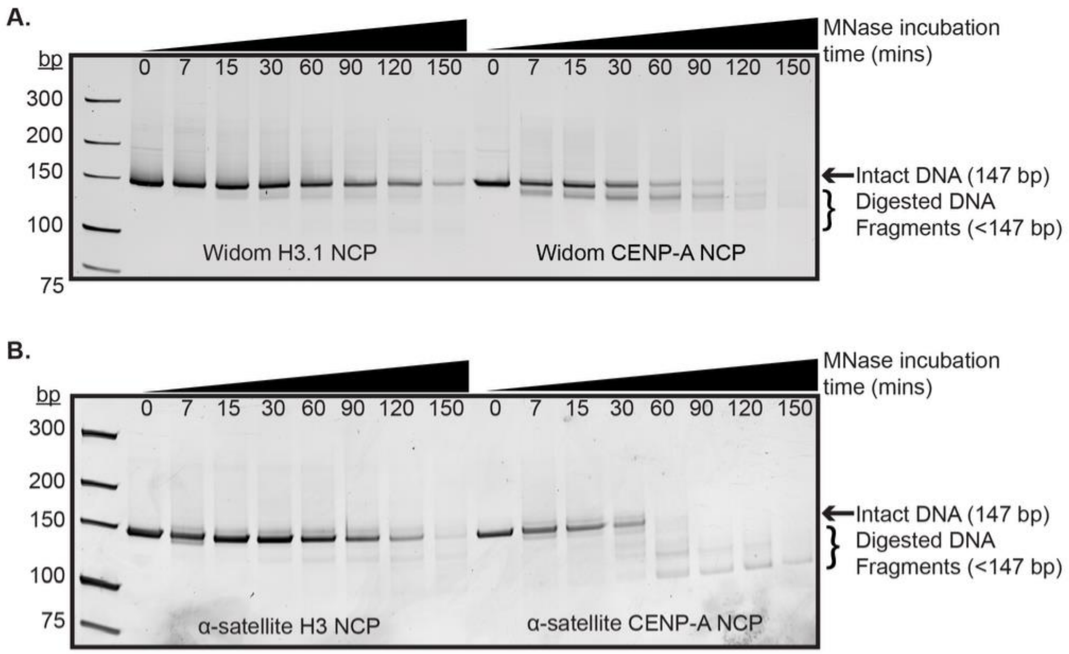


**Figure S5. WT NCP MNase Digestion. A.** Representative 10% TBE-Urea gel of Widom 601 H3 and CENP-A NCP MNase digestion. **B.** Representative 10% TBE-Urea gel of α-satellite H3 and CENP-A NCP MNase digestion.

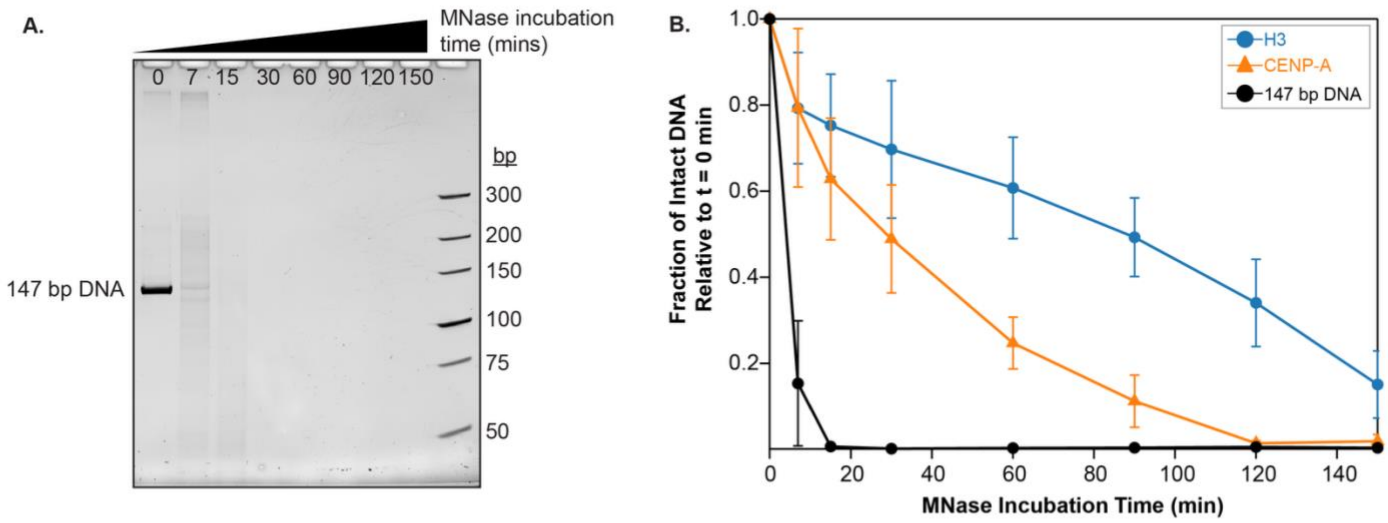


**Figure S6. Naked 147 bp 601 DNA MNase Digestion. A.** Representative MNase digestion 10% TBE-Urea gel. **B.** MNase digestion plot of H3 (blue circle), CENP-A (orange triangle), 147 bp DNA (black circle). Plots represent the average of a minimum of n=3 assays, and error bars represent the standard deviation of a minimum of n=3 assays.

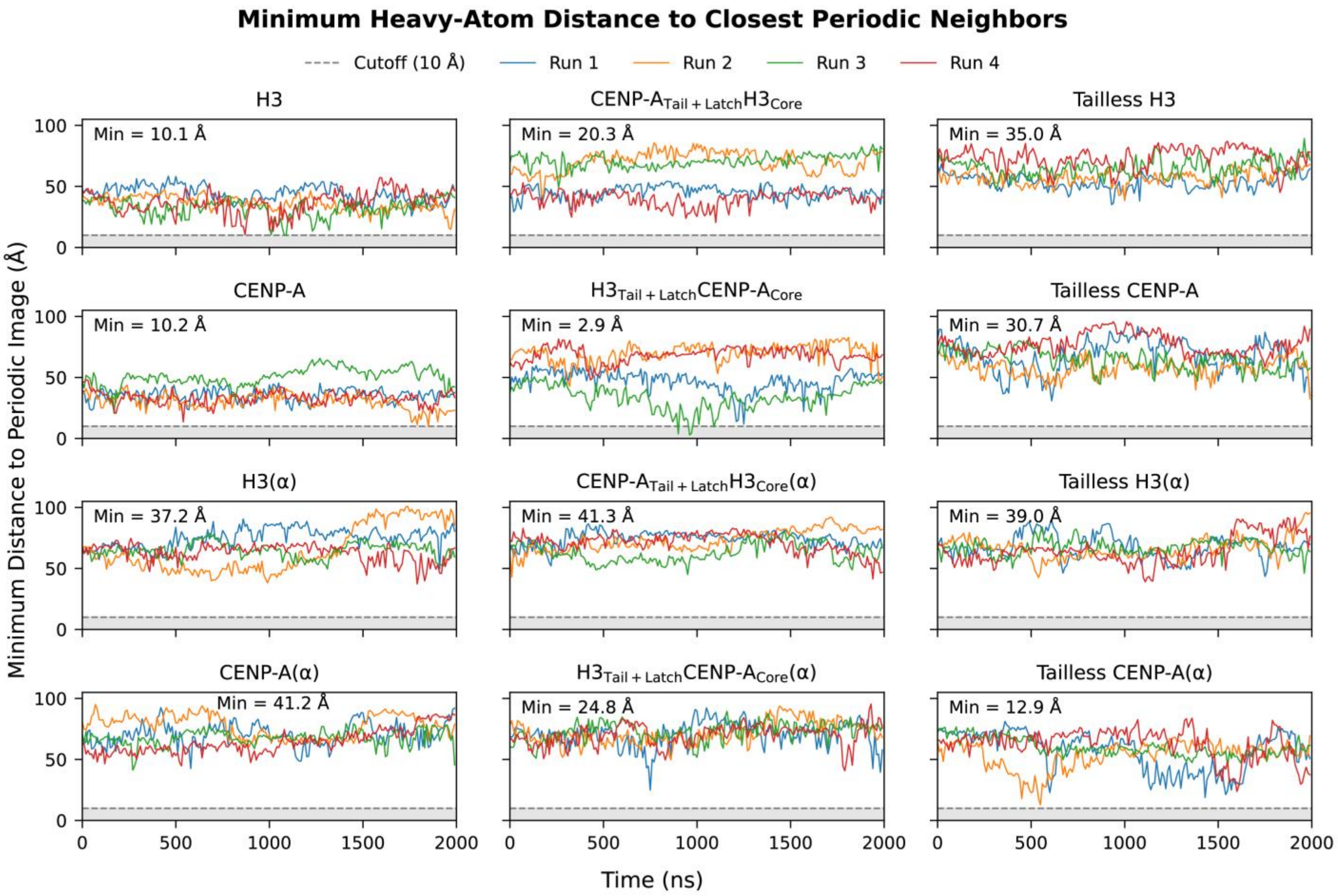


**Figure S7. Minimum Heavy-Atom Distances to Closest Periodic Neighbors in All-Atom NCPs.** Analysis of minimum distance between heavy atoms of the NCPs and their nearest periodic neighbor at any given moment in the trajectories. The dotted line above the gray area represents the 10 Angstrom electrostatic PME cutoff distance used. It is used to determine whether any long-term, consistent interactions between the NCP and any periodic neighbors occur. One instance of periodic contacts does occur in a replicate of H3$_{Tail+Latch}$CENP-A$_{Core}$, and accounts for 0.0417% of total all-atom frames across all NCPs.

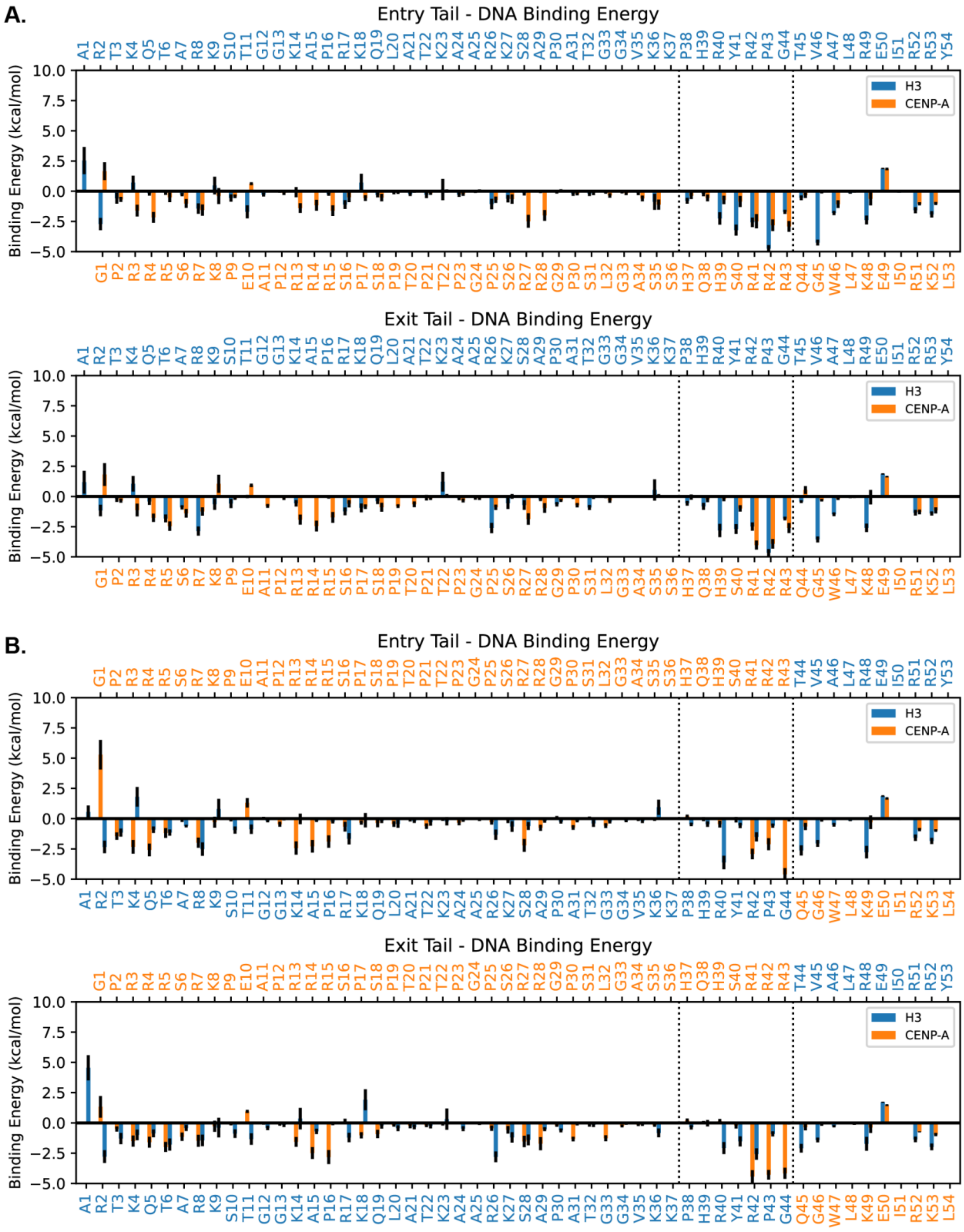


**Figure S8. MM/GBSA Per Residue Binding Energy Decomposition for All-Atom Widom 601 DNA NCPs. A.** All-atom Widom WT NCP entry and exit binding energy residue decomposition plots. **B.** All-atom Widom chimeric CENP-$A_{Tail+Latch}H3_{Core}$ and $H3_{Tail+Latch}CENP\text{-}A_{Core}$ NCP entry and exit binding energy residue decomposition plots. All energies are in kcal/mol. Error bars represent the standard error of the mean. In chimeric systems, both data bars and text labels are colored by histone of origin (H3 vs. CENP-A) in blue and orange, respectively.

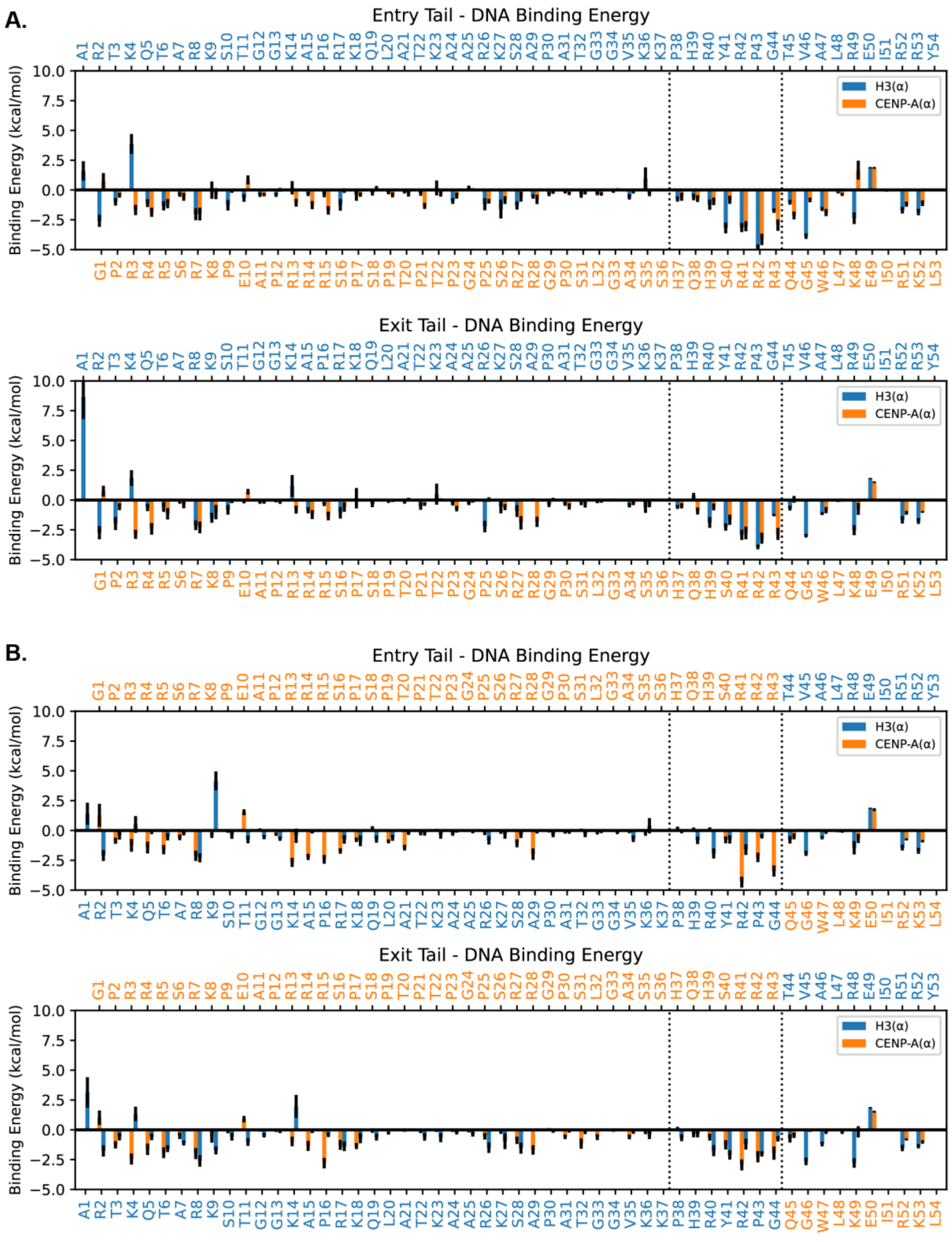


**Figure S9. MM/GBSA Per Residue Binding Energy Decomposition for All-Atom Alpha-Satellite DNA NCPs. A.** All-Atom alpha-satellite WT NCP entry and exit binding energy residue decomposition plots. **B.** All-atom alpha-satellite chimeric CENP-A$_{Tail+Latch}$H3$_{Core}$ and H3$_{Tail+Latch}$CENP-A$_{Core}$ NCP entry and exit binding energy residue decomposition plots. All energies are in kcal/mol. Error bars represent the standard error of the mean. In chimeric systems, both data bars and text labels are colored by histone of origin (H3 vs. CENP-A) in blue and orange, respectively.

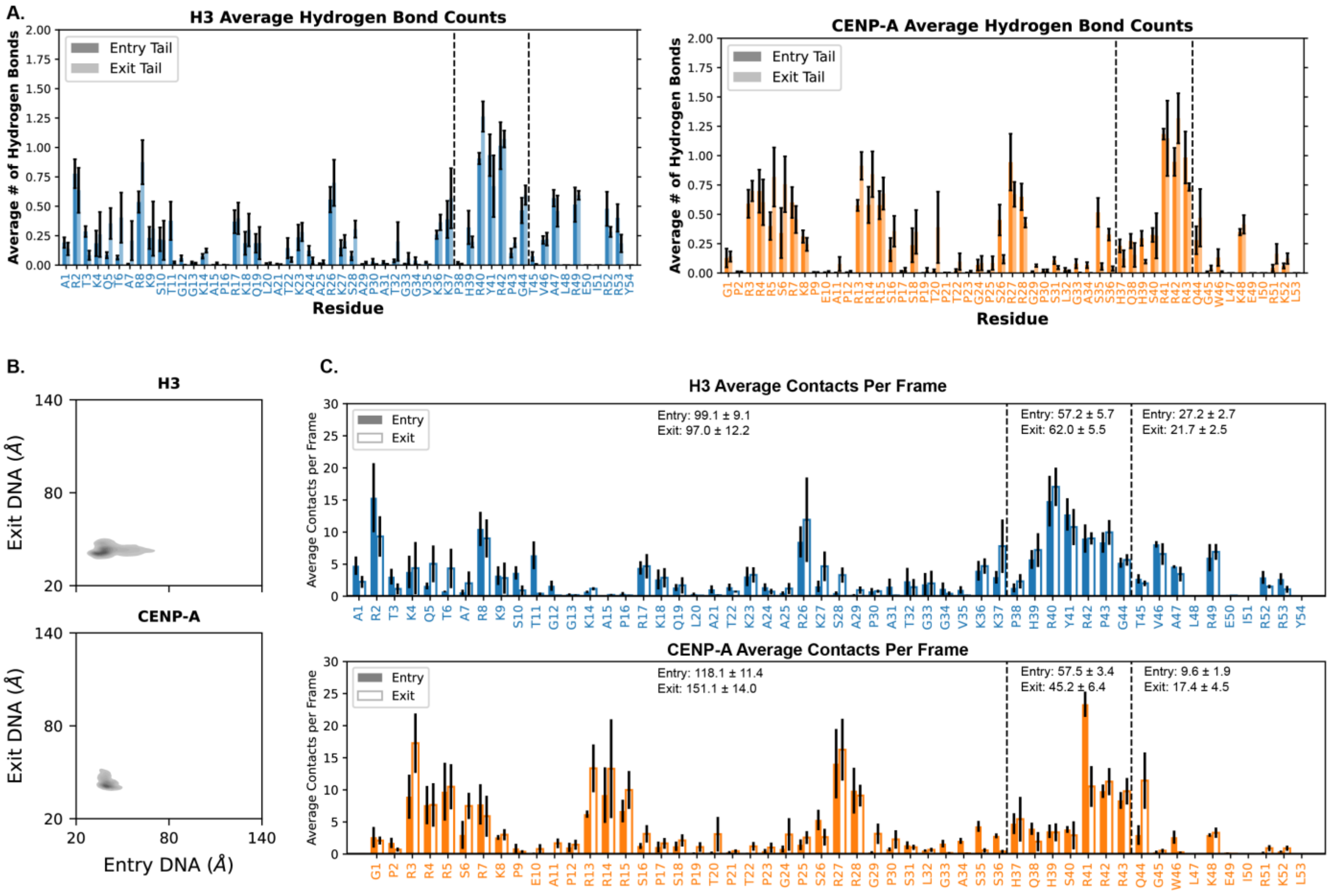


**Figure S10. All-Atom Widom 601 WT NCPs. A.** Per residue average hydrogen bonds for entry and exit N-terminal histone regions per nanosecond for all-atom Widom 601 WT NCPs. **B.** End-to-dyad plots for the terminal 5 bp of DNA of all-atom Widom 601 WT NCPs. **C.** Per residue average contacts per frame/nanosecond for all-atom Widom 601 WT NCPs. Entry tail contacts (solid bars) and Exit tail contacts (striped bars). Blue bars and text represent regions originating from H3; orange bars and text represent regions originating from CENP-A. Plots represent the average of n=4 replicas. Total average contacts for each region are above the data bars. All errors represent the standard error of the mean.

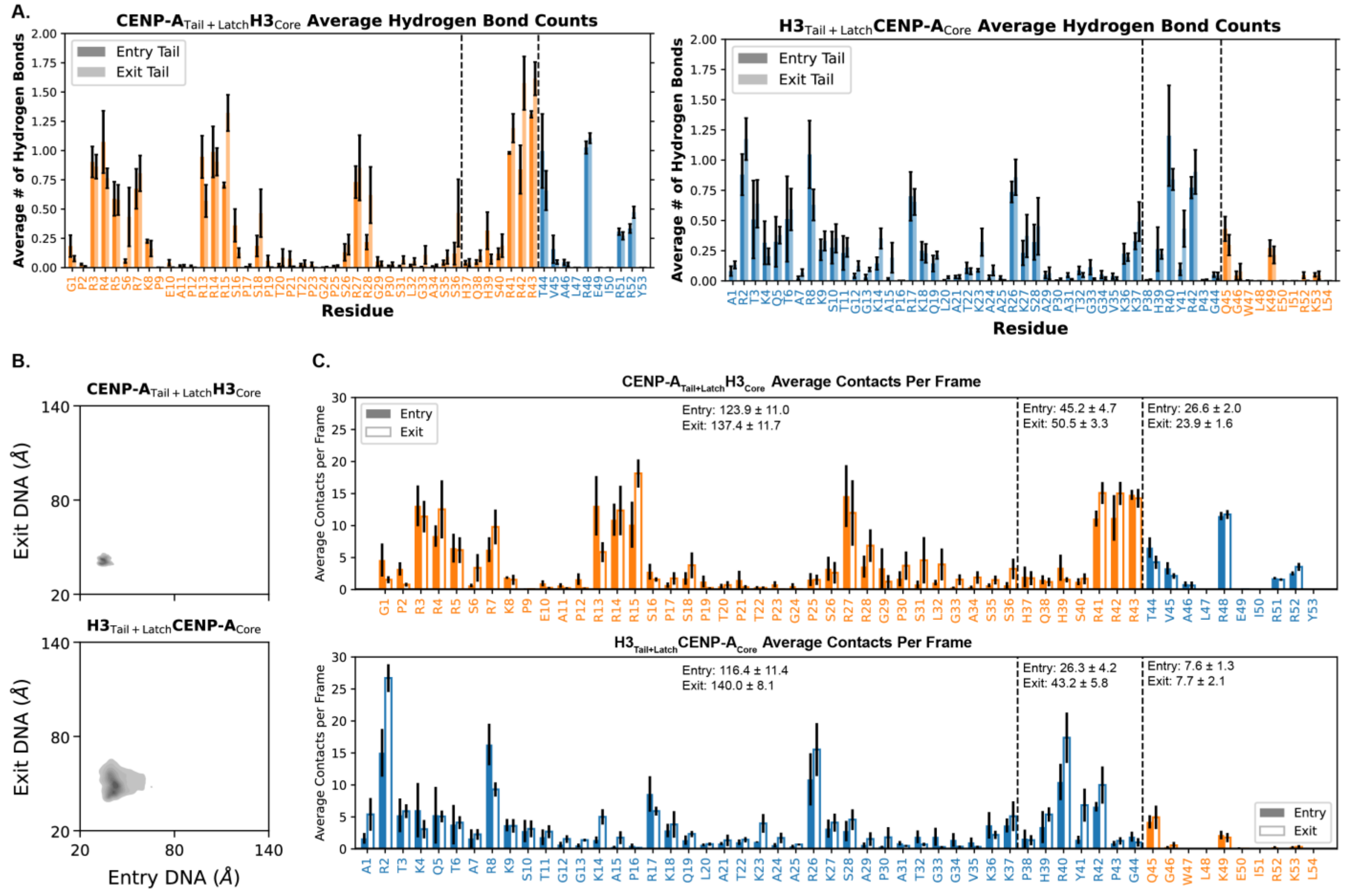


**Figure S11. All-Atom Widom 601 Chimera NCPs. A.** Per residue average hydrogen bonds for entry and exit N-terminal histone regions per nanosecond for all-atom Widom chimeric NCPs. **B.** End-to-dyad plots for the terminal 5 bp of DNA of all-atom Widom 601 chimeric NCPs. **C.** Per residue average contacts per frame/nanosecond for all-atom Widom 601 chimeric NCPs. Entry tail contacts (solid bars) and Exit tail contacts (striped bars). Blue bars and text represent regions originating from H3; orange bars and text represent regions originating from CENP-A. Plots represent the average of n=4 replicas. Total average contacts for each region are above the data bars. All errors represent the standard error of the mean.

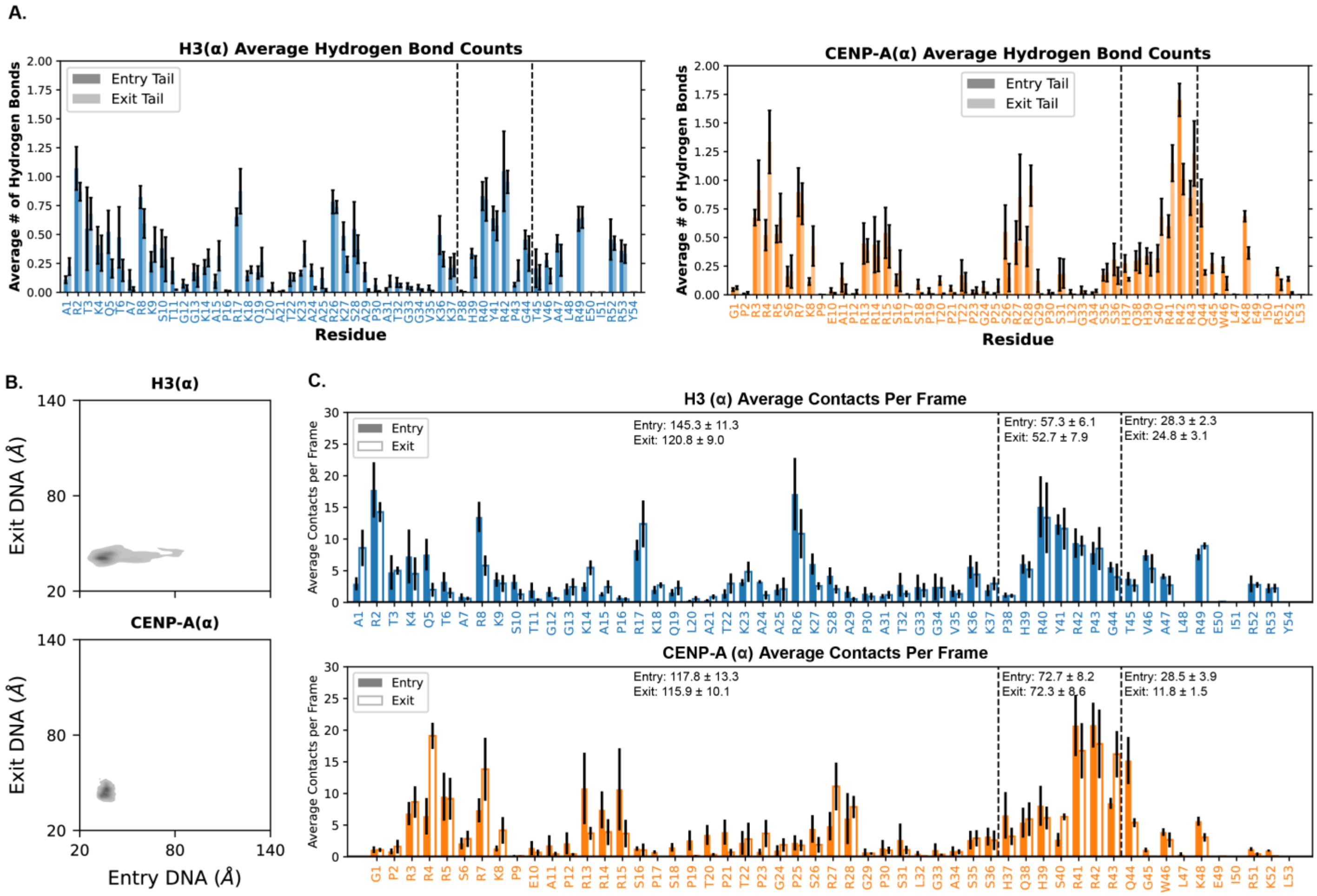


**Figure S12. All-Atom Alpha-Satellite WT NCPs. A.** Per residue average hydrogen bonds for entry and exit N-terminal histone regions per nanosecond for all-atom alpha-satellite WT NCPs. **B.** End-to-dyad plots for the terminal 5 bp of DNA of all-atom alpha-satellite WT NCPs. **C.** Per residue average contacts per frame/nanosecond for all-atom alpha-satellite WT NCPs. Entry tail contacts (solid bars) and Exit tail contacts (striped bars). Blue bars and text represent regions originating from H3; orange bars and text represent regions originating from CENP-A. Plots represent the average of n=4 replicas. Total average contacts for each region are above the data bars. All errors represent the standard error of the mean.

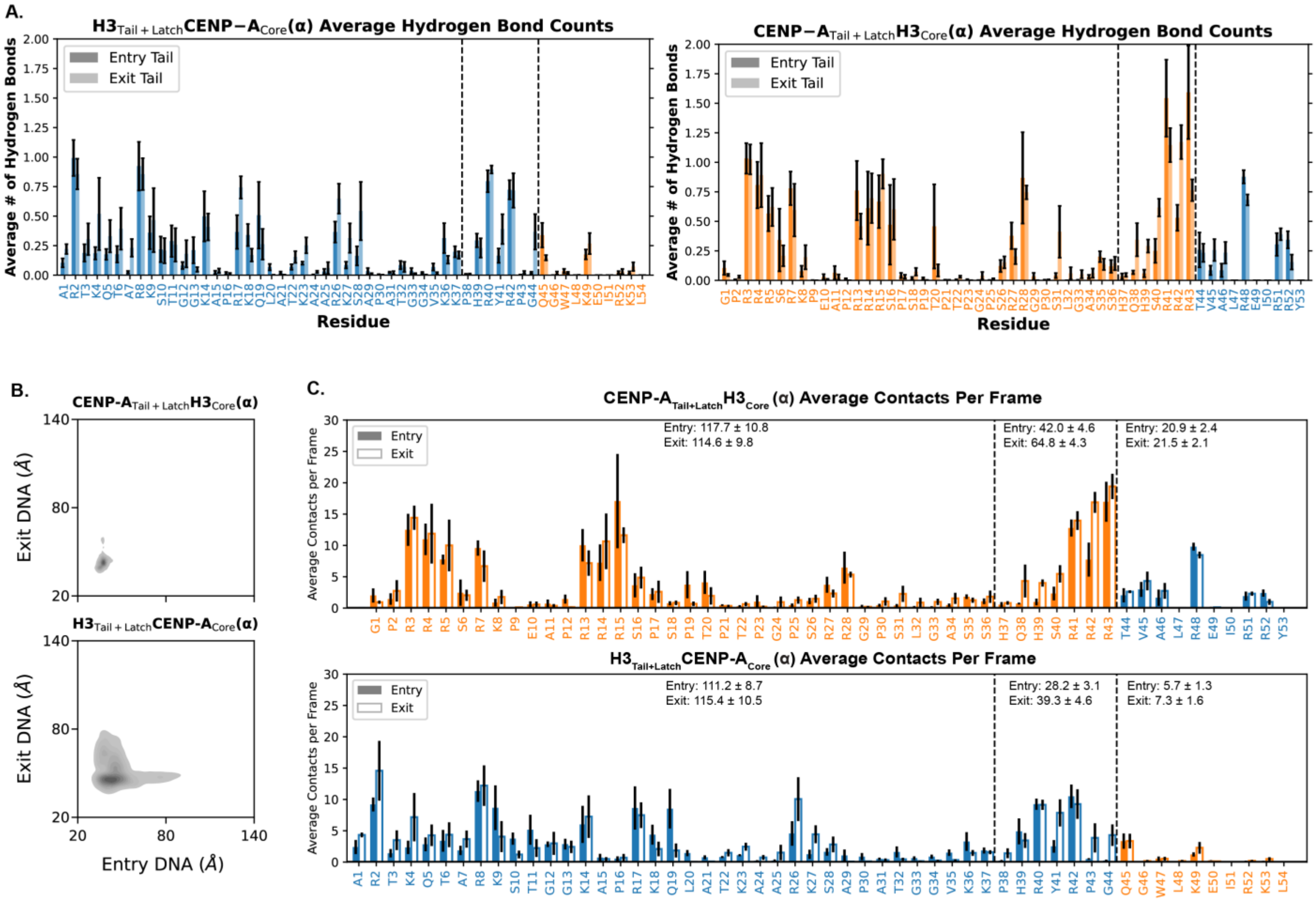


**Figure S13. All-Atom Alpha-Satellite Chimera NCPs. A.** Per residue average hydrogen bonds for entry and exit N-terminal histone regions per nanosecond for all-atom alpha-satellite chimeric NCPs. **B.** End-to-dyad plots for the terminal 5 bp of DNA of all-atom alpha-satellite chimeric NCPs. **C.** Per residue average contacts per frame/nanosecond for all-atom alpha-satellite chimeric NCPs. Entry tail contacts (solid bars) and Exit tail contacts (striped bars). Blue bars and text represent regions originating from H3; orange bars and text represent regions originating from CENP-A. Plots represent the average of n=4 replicas. Total average contacts for each region are above the data bars. All errors represent the standard error of the mean.

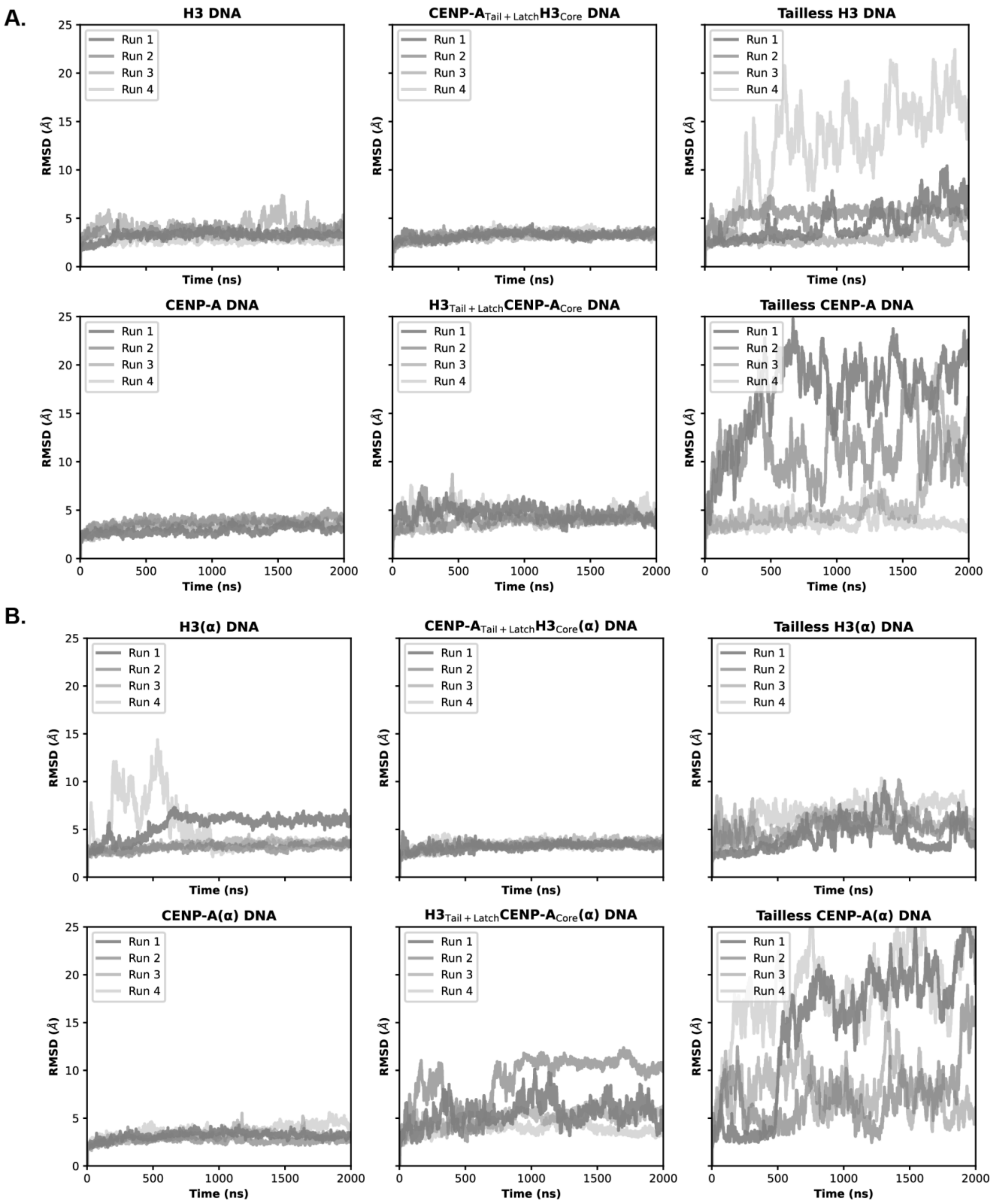


**Figure S14. All-Atom RMSD for All NCPs. A.** RMSD plots for all four replicas of each Widom DNA all-atom NCP. **B.** RMSD plots for all four replicas of each alpha-satellite DNA all-atom NCP. Alignment of all RMSD calculations was to the first frame histone H4 pair core residues for each respective trajectory for consistency across all NCPs. Minimal differences were observed between identical systems across both DNA sequences.

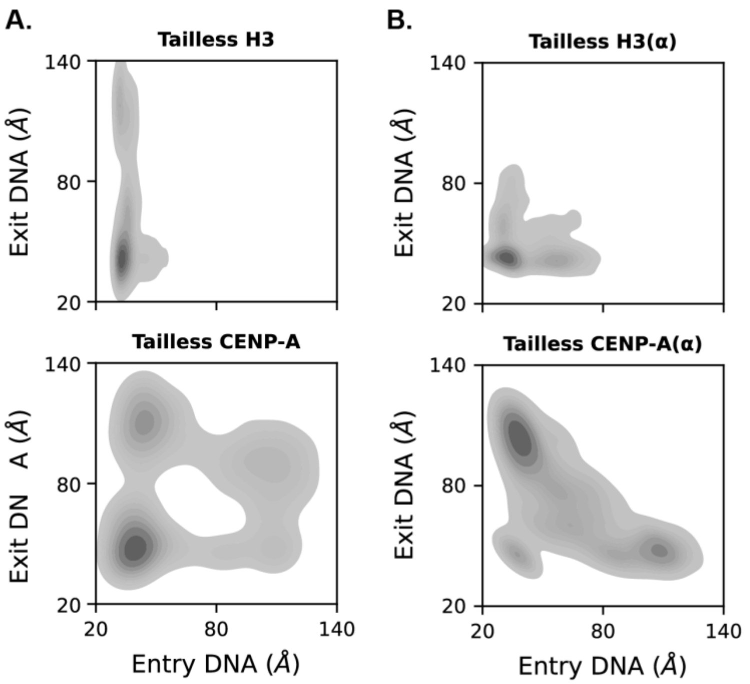


**Figure S15. All-Atom End-to-Dyad of Tailless NCPs. A.** End-to-dyad plots for the terminal 5 bp of DNA of all-atom Widom DNA tailless NCPs. **B.** End-to-dyad plots for the terminal 5 bp of DNA of all-atom alpha-satellite DNA tailless NCPs. Increased DNA accessibility observed in all cases compared to WTs.

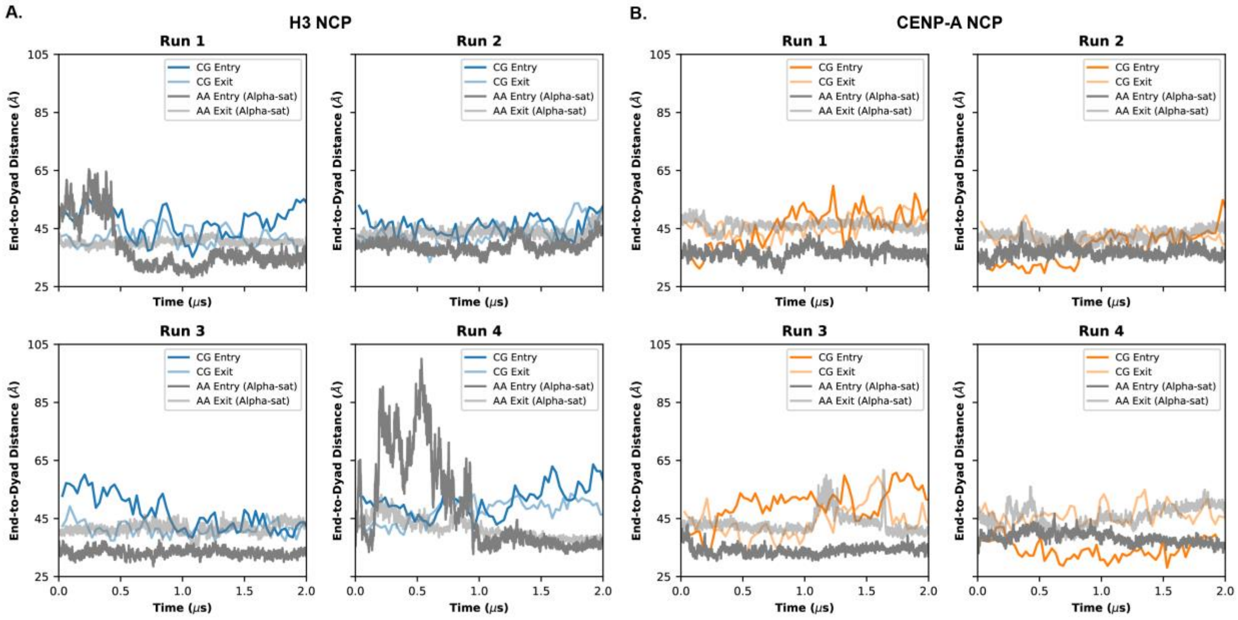


**Figure S16. All-Atom and Coarse-Grained NCPs End-to-Dyad Distances Over Comparable Timescales. A.** Time-resolved end-to-dyad plots comparing the all-atom H3 with alpha-satellite DNA to the coarse-grained H3 with alpha-satellite DNA on a roughly equivalent timescale. The CG data is zoomed into the initial steps calculated to be the estimated equivalent of 2 μs, which is the full length of the all-atom simulations. **B.** Time-resolved end-to-dyad plots comparing the all-atom CENP-A with alpha-satellite DNA to the coarse-grained CENP-A with alpha-satellite DNA on a roughly equivalent timescale. The CG data is zoomed into the initial steps calculated to be the estimated equivalent of 2 μs, which is the full length of the all-atom simulations. Both CG NCPs show comparable end-to-dyad distances for both entry and exit as their all-atom equivalents using the same alpha-satellite DNA sequence.

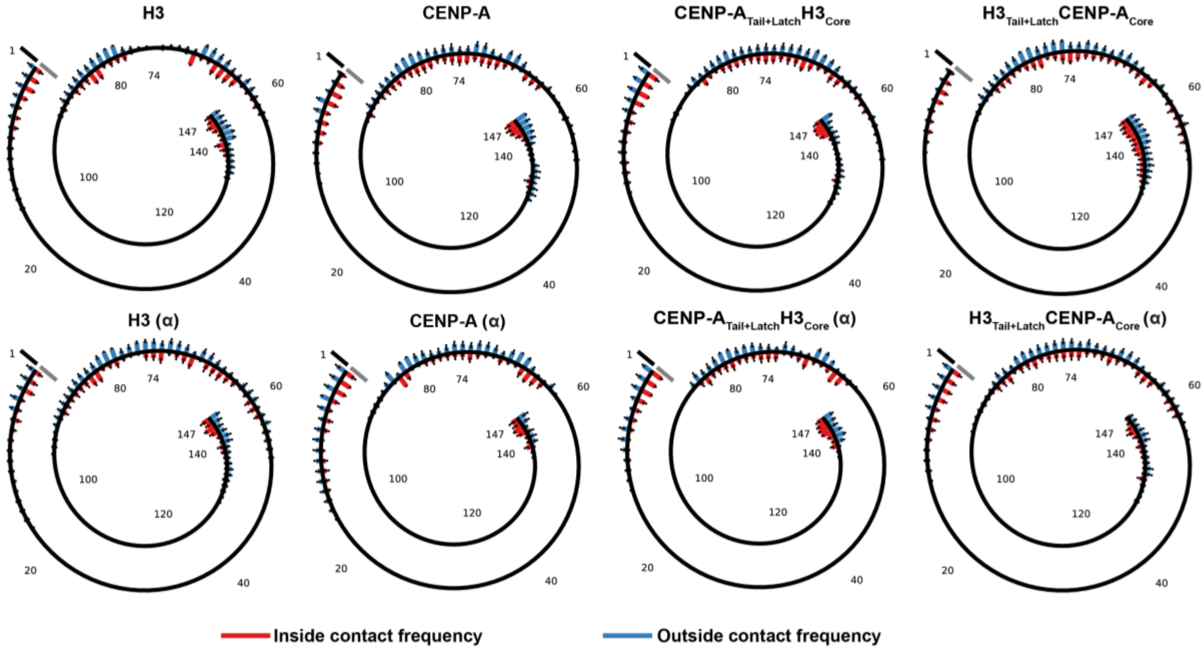


**Figure S17. Tail-DNA Contacts Positioning and Orientations Along the DNA for All-Atom NCPs.** Spiral plots showing tail contact positioning along the DNA with base pairs labeled. NCPs on the top have Widom 601 DNA. NCPs on the bottom have centromeric alpha-satellite DNA. Inside and outside contacts represented by red and blue bars, respectively. Black and gray reference bars represent the maximum frequency of contacts able to be made per NCP type, with 0 meaning no contacts made throughout the trajectories and 1 meaning a tail-DNA contact was persistent across the entire trajectory. All error bars represent the standard error of the mean.

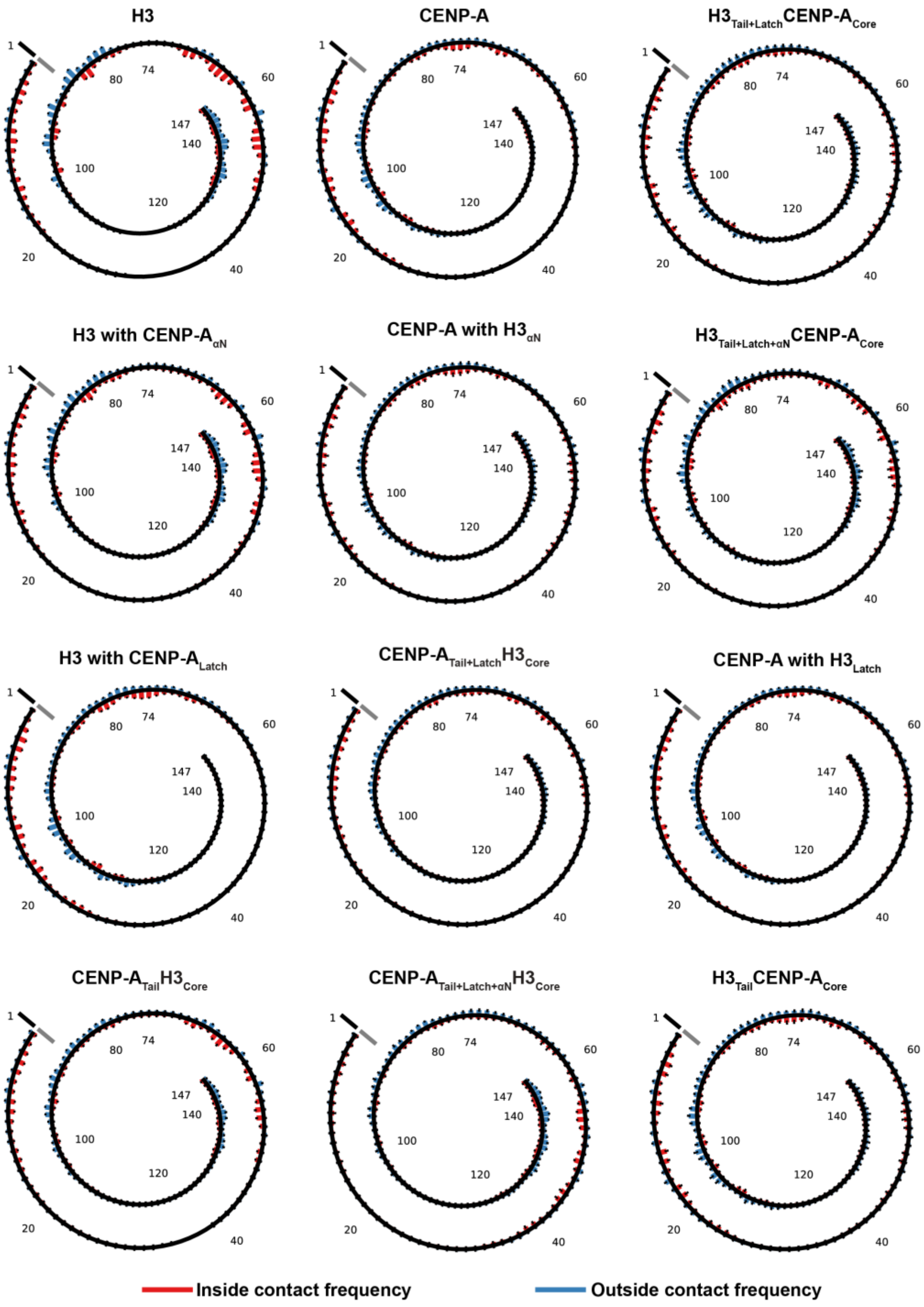


**Figure S18. Tail-DNA Contacts Positioning and Orientations Along the DNA for Coarse-Grained NCPs.** Spiral plots showing tail contact positioning along the DNA with base pairs labeled. Left half are NCPs that have the histone H3 core (below the αN-helix); right half are NCPs that have the histone CENP-A core (below the αN-helix). Inside and outside contacts represented by red and blue bars, respectively. Black and gray reference bars represent the maximum frequency of contacts able to be made per NCP type, with 0 meaning no contacts made throughout the trajectories and 1 meaning a tail-DNA contact was persistent across the entire trajectory. All error bars represent the standard error of the mean.

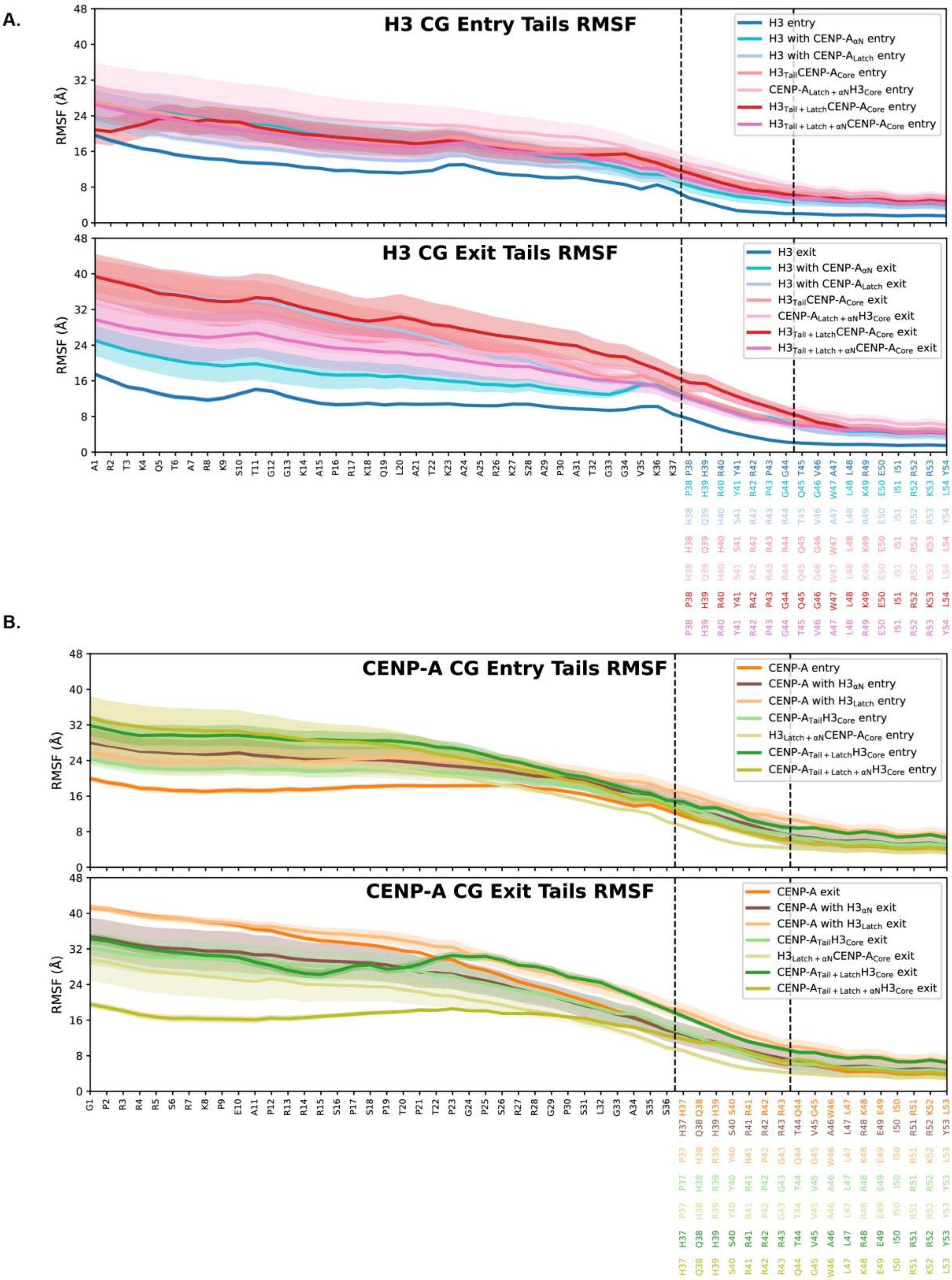


**Figure S19. Coarse-Grained RMSF of H3/CENP-A N-terminal Histone Regions for all Tailed NCPs. A.** RMSF of NCPs that share the H3 tail residues (1-37). Entry RMSFs depicted on top, exit RMSFs on bottom. **B.** RMSF of NCPs that share the CENP-A tail residues (1-36). Entry RMSFs depicted on top, exit RMSFs on bottom. Alignment of all RMSF calculations was to the first frame histone H4 pair core residues for each respective trajectory for consistency across all NCPs. Lines represent the mean values. Error spreads represent the standard error of the mean.

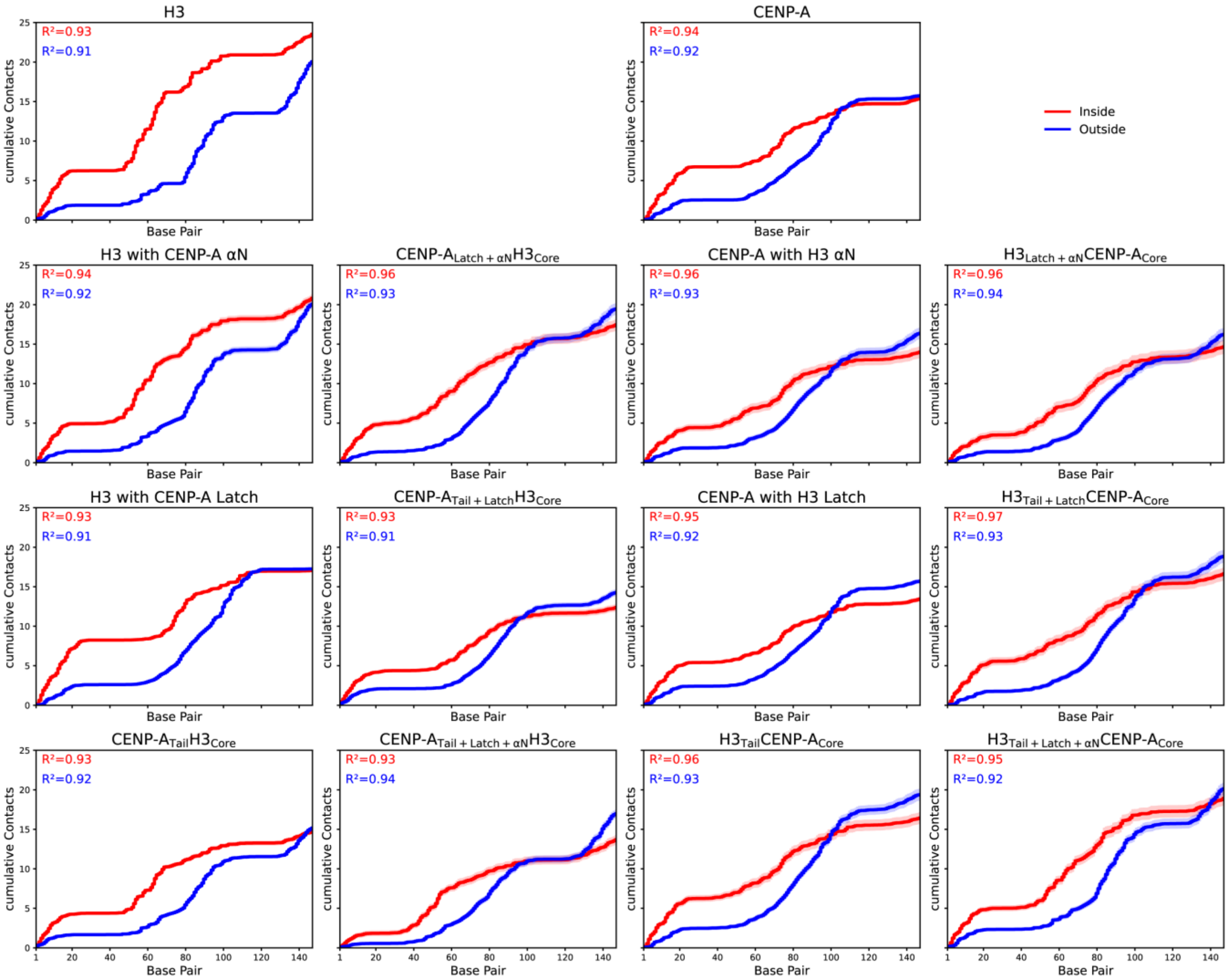


**Figure S20. CG Simulation Cumulative Contacts Between H3/CENP-A N-terminal Tails and DNA Across All DNA Base Pairs.** Stepwise cumulative contacts plots between histone tails and DNA representing contacts made along the nucleosomal DNA. Increases in the data lines mean contacts are being made. Flat areas indicate regions along the DNA where the tails and DNA do contact. $R^2$ values provided as a measure of how consistent contacts are being made along the DNA for contact orientation (inside vs. outside). Red lines represent average inside contacts being made. Blue lines represent average outside contacts being made. Error spreads represent the standard error of the mean.

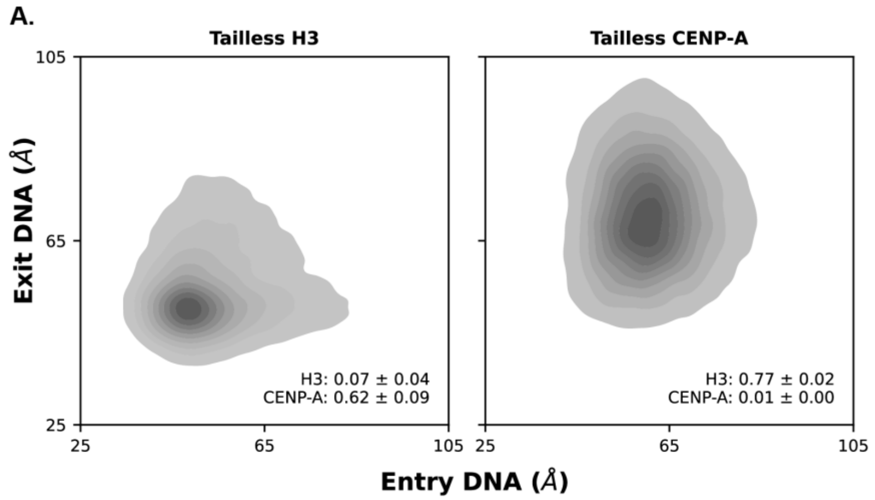


**Figure S21. Coarse-Grained Tailless Systems End-to-Dyad.** End-to-dyad plots for the terminal 5 bp of DNA of CG tailless NCPs. Increased DNA accessibility observed compared to WTs.

### Supplemental Tables.

| System | # of Replicas | Equilibration Time (per copy) | Simulation Time (per copy) | Simulation Time (Total) |
|---|---|---|---|---|
| H3 | 4 | 300 ns | 2000 ns | 2300 ns |
| CENP-A | | | | |
| CENP-A$_{Tail+Latch}$H3$_{Core}$ | | | | |
| H3$_{Tail+Latch}$CENP-A$_{Core}$ | | | | |
| Tailless H3 | | | | |
| Tailless CENP-A | | | | |

**Table S1. All-Atom Simulation Time Details.** All simulations were run for the same amount of time. The values given in this table are applicable to all all-atom simulations regardless of which DNA sequence was used.

| System | # of Replicas | Equilibration Time (per copy) | Simulation Time (per copy) | Simulation Time (Total) |
|---|---|---|---|---|
| H3 | 4 | 600 μs | 2400 μs | 3000 μs |
| CENP-A | | | | |
| CENP-A$_{Tail+Latch+\alpha N}$H3$_{Core}$ | | | | |
| H3$_{Tail+Latch+\alpha N}$CENP-A$_{Core}$ | | | | |
| CENP-A$_{Tail+Latch}$H3$_{Core}$ | | | | |
| H3$_{Tail+Latch}$CENP-A$_{Core}$ | | | | |
| CENP-A$_{Latch+\alpha N}$H3$_{Core}$ | | | | |
| H3$_{Latch+\alpha N}$CENP-A$_{Core}$ | | | | |
| CENP-A$_{Tail}$H3$_{Core}$ | | | | |
| H3$_{Tail}$CENP-A$_{Core}$ | | | | |
| H3 with CENP-A$_{Latch}$ | | | | |
| CENP-A with H3$_{Latch}$ | | | | |
| H3 with CENP-A$_{\alpha N}$ | | | | |
| CENP-A with H3$_{\alpha N}$ | | | | |
| Tailless H3 | | | | |
| Tailless CENP-A | | | | |

**Table S2. Coarse-Grained Simulation Time Details.** All CG simulations were run for the same amount of time.

| Widom Entry Tails | | | |
|---|---|---|---|
| System | Δvdw (kcal/mol) | Δelec (kcal/mol) | ΔTotal (kcal/mol) |
| H3 | -67.3 ± 6.7 | 10.0 ± 10.6 | -57.4 ± 12.6 |
| CENP-A | -76.4 ± 3.7 | -1.7 ± 5.9 | -78.2 ± 6.9 |
| CENP-$A_{Tail+Latch}H3_{Core}$ | -82.2 ± 3.3 | 4.3 ± 5.8 | -78.0 ± 6.7 |
| $H3_{Tail+Latch}$CENP-$A_{Core}$ | -77.3 ± 5.0 | 3.9 ± 8.8 | -73.4 ± 10.1 |
| Widom Exit Tails | | | |
| System | Δvdw (kcal/mol) | Δelec (kcal/mol) | ΔTotal (kcal/mol) |
| H3 | -65.0 ± 3.8 | -0.1 ± 5.6 | -65.2 ± 6.8 |
| CENP-A | -100.6 ± 5.1 | 10.9 ± 8.4 | -89.8 ± 9.9 |
| CENP-$A_{Tail+Latch}H3_{Core}$ | -90.6 ± 5.2 | 1.3 ± 8.7 | -89.3 ± 10.1 |
| $H3_{Tail+Latch}$CENP-$A_{Core}$ | -95.5 ± 3.9 | 6.8 ± 6.6 | -88.7 ± 7.6 |

**Table S3. Widom Tails MM/GBSA Energetics Values.** Entry and exit values given in kcal/mol. Errors represent the standard error of the mean.

| Widom Entry Latch | | | |
|---|---|---|---|
| System | Δvdw (kcal/mol) | Δelec (kcal/mol) | ΔTotal (kcal/mol) |
| H3 | -44.4 ± 6.3 | 6.1 ± 9.1 | -38.3 ± 11.1 |
| CENP-A | -38.8 ± 2.4 | 1.6 ± 4.2 | -37.2 ± 4.8 |
| CENP-$A_{Tail+Latch}H3_{Core}$ | -30.1 ± 6.2 | -3.1 ± 10.7 | -33.2 ± 12.3 |
| $H3_{Tail+Latch}$CENP-$A_{Core}$ | -17.5 ± 3.7 | -1.6 ± 6.5 | -19.1 ± 7.5 |
| Widom Exit Latch | | | |
| System | Δvdw (kcal/mol) | Δelec (kcal/mol) | ΔTotal (kcal/mol) |
| H3 | -51.1 ± 3.6 | 10.5 ± 5.2 | -40.6 ± 6.4 |
| CENP-A | -29.2 ± 2.7 | -4.4 ± 4.5 | -33.6 ± 5.3 |
| CENP-$A_{Tail+Latch}H3_{Core}$ | -30.6 ± 4.4 | -9.1 ± 7.5 | -39.7 ± 8.7 |
| $H3_{Tail+Latch}$CENP-$A_{Core}$ | -32.7 ± 4.7 | 8.3 ± 8.2 | -24.4 ± 9.4 |

**Table S4. Widom Latch MM/GBSA Energetics Values.** Entry and exit values given in kcal/mol. Errors represent the standard error of the mean.

| Widom Entry αN | | | |
|---|---|---|---|
| System | Δvdw (kcal/mol) | Δelec (kcal/mol) | ΔTotal (kcal/mol) |
| H3 | -19.1 ± 4.6 | -3.1 ± 6.9 | -22.3 ± 8.3 |
| CENP-A | -11.6 ± 2.6 | 2.9 ± 4.5 | -8.6 ± 5.2 |
| CENP-$A_{Tail+Latch}H3_{Core}$ | -18.2 ± 5.9 | -4.7 ± 10.1 | -23.0 ± 11.7 |
| $H3_{Tail+Latch}$CENP-$A_{Core}$ | -8.1 ± 4.1 | 1.7 ± 7.1 | -6.4 ± 8.2 |
| Tailless H3 | -18.9 ± 8.3 | -0.9 ± 13.6 | -19.9 ± 15.9 |
| Tailless CENP-A | -12.1 ± 4.6 | 4.1 ± 7.8 | -8.0 ± 9.0 |
| Widom Exit αN | | | |
| System | Δvdw (kcal/mol) | Δelec (kcal/mol) | ΔTotal (kcal/mol) |
| H3 | -16.2 ± 2.8 | -1.8 ± 4.3 | -18.0 ± 5.1 |
| CENP-A | -17.9 ± 3.8 | 8.2 ± 6.4 | -9.6 ± 7.4 |
| CENP-$A_{Tail+Latch}H3_{Core}$ | -17.2 ± 7.3 | -2.7 ± 12.5 | -19.8 ± 14.5 |
| $H3_{Tail+Latch}$CENP-$A_{Core}$ | -8.4 ± 3.9 | 2.6 ± 6.9 | -5.8 ± 7.9 |
| Tailless H3 | -15.4 ± 7.8 | 0.2 ± 13.0 | -15.2 ± 15.1 |
| Tailless CENP-A | -6.4 ± 4.3 | 2.1 ± 7.3 | -4.3 ± 8.4 |

**Table S5. Widom αN-Helix MM/GBSA Energetics Values.** Entry and exit values given in kcal/mol. Errors represent the standard error of the mean.

| Alpha-sat Entry Tails | | | |
|---|---|---|---|
| System | Δvdw (kcal/mol) | Δelec (kcal/mol) | ΔTotal (kcal/mol) |
| H3 | -99.9 ± 7.1 | 11.7 ± 12.3 | -88.2 ± 14.2 |
| CENP-A | -80.9 ± 5.1 | 10.4 ± 8.0 | -70.6 ± 9.5 |
| CENP-$A_{Tail+Latch}H3_{Core}$ | -76.4 ± 5.6 | -3.5 ± 8.8 | -80.0 ± 10.5 |
| $H3_{Tail+Latch}$CENP-$A_{Core}$ | -78.5 ± 7.5 | 14.6 ± 11.9 | -63.9 ± 14.1 |
| Alpha-sat Exit Tails | | | |
| System | Δvdw (kcal/mol) | Δelec (kcal/mol) | ΔTotal (kcal/mol) |
| H3 | -80.1 ± 3.3 | 4.1 ± 5.3 | -76.0 ± 6.3 |
| CENP-A | -74.0 ± 7.6 | -2.1 ± 11.7 | -76.2 ± 13.9 |
| CENP-$A_{Tail+Latch}H3_{Core}$ | -75.9 ± 4.7 | -3.3 ± 7.4 | -79.2 ± 8.7 |
| $H3_{Tail+Latch}$CENP-$A_{Core}$ | -78.8 ± 9.5 | -0.4 ± 15.1 | -79.2 ± 17.9 |

**Table S6. Alpha-Satellite Tails MM/GBSA Energetics Values.** Entry and exit values given in kcal/mol. Errors represent the standard error of the mean.

| Alpha-sat Entry Latch | | | |
|---|---|---|---|
| System | Δvdw (kcal/mol) | Δelec (kcal/mol) | ΔTotal (kcal/mol) |
| H3 | -47.2 ± 5.2 | 7.8 ± 8.8 | -39.4 ± 10.2 |
| CENP-A | -50.2 ± 3.4 | 6.1 ± 5.5 | -44.1 ± 6.5 |
| CENP-A$_{Tail+Latch}$H3$_{Core}$ | -26.7 ± 7.7 | -5.4 ± 12.0 | -32.2 ± 14.2 |
| H3$_{Tail+Latch}$CENP-A$_{Core}$ | -20.5 ± 6.1 | 2.7 ± 10.1 | -17.8 ± 11.8 |
| Alpha-sat Exit Latch | | | |
| System | Δvdw (kcal/mol) | Δelec (kcal/mol) | ΔTotal (kcal/mol) |
| H3 | -40.4 ± 6.7 | 7.0 ± 11.2 | -33.3 ± 13.1 |
| CENP-A | -48.3 ± 5.4 | 4.6 ± 9.1 | -43.6 ± 10.6 |
| CENP-A$_{Tail+Latch}$H3$_{Core}$ | -44.3 ± 2.1 | 6.2 ± 3.4 | -38.1 ± 4.0 |
| H3$_{Tail+Latch}$CENP-A$_{Core}$ | -29.9 ± 5.1 | 4.4 ± 8.5 | -25.5 ± 9.9 |

**Table S7. Alpha-Satellite Latch MM/GBSA Energetics Values.** Entry and exit values given in kcal/mol. Errors represent the standard error of the mean.

| Alpha-sat Entry αN | | | |
|---|---|---|---|
| System | Δvdw (kcal/mol) | Δelec (kcal/mol) | ΔTotal (kcal/mol) |
| H3 | -19.6 ± 4.2 | -2.6 ± 7.4 | -22.2 ± 8.5 |
| CENP-A | -25.8 ± 4.6 | 7.4 ± 7.7 | -18.4 ± 8.9 |
| CENP-A$_{Tail+Latch}$H3$_{Core}$ | -17.7 ± 4.3 | 1.3 ± 6.9 | -16.4 ± 8.1 |
| H3$_{Tail+Latch}$CENP-A$_{Core}$ | -7.0 ± 5.7 | 2.0 ± 9.5 | -5.0 ± 11.1 |
| Tailless H3 | -20.2 ± 6.7 | 3.2 ± 10.5 | -17.0 ± 12.5 |
| Tailless CENP-A | -12.4 ± 1.7 | 5.8 ± 2.8 | -6.5 ± 3.3 |
| Alpha-sat Exit αN | | | |
| System | Δvdw (kcal/mol) | Δelec (kcal/mol) | ΔTotal (kcal/mol) |
| H3 | -18.2 ± 4.8 | -1.6 ± 8.3 | -19.8 ± 9.6 |
| CENP-A | -14.5 ± 3.3 | 5.1 ± 5.5 | -9.4 ± 6.4 |
| CENP-A$_{Tail+Latch}$H3$_{Core}$ | -17.2 ± 4.3 | -1.5 ± 7.0 | -18.7 ± 8.2 |
| H3$_{Tail+Latch}$CENP-A$_{Core}$ | -10.0 ± 5.4 | 3.5 ± 9.2 | -6.5 ± 10.6 |
| Tailless H3 | -18.1 ± 8.1 | -1.8 ± 12.6 | -19.9 ± 15.0 |
| Tailless CENP-A | -8.7 ± 2.3 | 2.4 ± 4.0 | -6.2 ± 4.7 |

**Table S8. Alpha-Satellite αN-Helix MM/GBSA Energetics Values.** Entry and exit values given in kcal/mol. Errors represent the standard error of the mean.

| System | H3 KL Divergence | H3 KL Std Err of Mean | CENP-A KL Divergence | CENP-A KL Std Err of Mean |
|---|---|---|---|---|
| H3 | 0.00 | 0.00 | 1.00 | 0.01 |
| CENP-A | 0.79 | 0.04 | 0.00 | 0.00 |
| CENP-A$_{Tail+Latch+\alpha N}$H3$_{Core}$ | 0.60 | 0.22 | 0.10 | 0.07 |
| H3$_{Tail+Latch+\alpha N}$CENP-A$_{Core}$ | 0.28 | 0.40 | 0.29 | 0.32 |
| CENP-A$_{Tail+Latch}$H3$_{Core}$ | 0.47 | 0.23 | 0.13 | 0.13 |
| H3$_{Tail+Latch}$CENP-A$_{Core}$ | 0.84 | 0.25 | 0.03 | 0.02 |
| CENP-A$_{Latch+\alpha N}$H3$_{Core}$ | 0.57 | 0.25 | 0.05 | 0.09 |
| H3$_{Latch+\alpha N}$CENP-A$_{Core}$ | 0.76 | 0.24 | 0.01 | 0.04 |
| CENP-A$_{Tail}$H3$_{Core}$ | 0.19 | 0.10 | 0.41 | 0.15 |
| H3$_{Tail}$CENP-A$_{Core}$ | 0.75 | 0.27 | 0.02 | 0.08 |
| H3 with CENP-A$_{Latch}$ | 1.00 | 0.02 | 0.04 | 0.01 |
| CENP-A with H3$_{Latch}$ | 0.99 | 0.08 | 0.03 | 0.02 |
| H3 with CENP-A$_{\alpha N}$ | 0.21 | 0.11 | 0.30 | 0.14 |
| CENP-A with H3$_{\alpha N}$ | 0.86 | 0.22 | 0.01 | 0.02 |
| Tailless H3 | 0.07 | 0.04 | 0.62 | 0.09 |
| Tailless CENP-A | 0.77 | 0.02 | 0.01 | 0.00 |

**Table S9. Kullback-Leibler Divergence Values from CG End-to-Dyad Data.** KL values for all NCPs for both H3-based and CENP-A-based calculations are provided with errors representing the standard error of the mean. Widom 601 DNA coarse-grained NCPs represent the last four rows, denominated by the (601) in the System titles. All calculations are based on the H3 or CENP-A alpha-satellite CG NCPs.

| System | Solvation buffer (Å) | No. of water molecules | Initial box XYZ dimensions (Å) | Initial box volume ($Å^3$) |
|---|---|---|---|---|
| H3 | 16x16x16 | 123,493 | 149.35x268.94x112.38 | 4,514,093.79 |
| CENP-A | 16x16x16 | 107,865 | 145.65x259.79x106.53 | 4,030,731.85 |
| CENP-$A_{Tail+Latch}$H$3_{Core}$ | 16x16x16 | 118,051 | 149.35x259.51x112.38 | 4,355,893.74 |
| H$3_{Tail+Latch}$CENP-$A_{Core}$ | 16x16x16 | 115,324 | 145.65x271.75x108.45 | 4,292,519.96 |
| Tailless H3 | 18x18x18 | 70,063 | 153.07x154.06x116.67 | 2,751,452.60 |
| Tailless CENP-A | 18x18x18 | 63,954 | 149.66x154.35x110.83 | 2,560,122.89 |

**Table S10. All-Atom Widom 601 DNA Equilibration Water Box Dimensions.** Water box solvation buffers, number of water molecules, and initial box dimensions for all-atom Widom 601 NCPs during equilibration phase (300 ns). Solvation buffer is the specified distance. Initial box dimensions are the actual water box XYZ dimensions which encase the all-atom nucleosome structures.

| System | Solvation buffer (Å) | No. of water molecules | Initial box XYZ dimensions (Å) | Initial box volume ($Å^3$) |
|---|---|---|---|---|
| H3 | 20x20x20 | 145,744 | 156.84x277.17x120.89 | 5,255,221.95 |
| CENP-A | 20x20x20 | 129,251 | 153.41x267.87x114.92 | 4,722,166.49 |
| CENP-$A_{Tail+Latch}$H$3_{Core}$ | 30x30x30 | 205,053 | 177.07x287.39x140.75 | 7,162,177.03 |
| H$3_{Tail+Latch}$CENP-$A_{Core}$ | 30x30x30 | 203,176 | 173.80x299.62x136.24 | 7,094,456.10 |
| Tailless H3 | 50x30x50 | 200,152 | 216.98x176.79x180.80 | 6,935,227.94 |
| Tailless CENP-A | 50x30x50 | 191,052 | 213.72x178.42x174.75 | 6,663,715.42 |

**Table S11. All-Atom Alpha-Satellite DNA Equilibration Water Box Dimensions.** Water box solvation buffers, number of water molecules, and initial box dimensions for all-atom alpha-satellite NCPs during equilibration phase (300 ns). Solvation buffer is the specified distance. Initial box dimensions are the actual water box XYZ dimensions which encase the all-atom nucleosome structures.

| System | Run # | Solvation buffer (Å) | No. of water molecules | Initial box XYZ dimensions (Å) | Initial box volume ($Å^3$) |
|---|---|---|---|---|---|
| H3 | 1 | 20x20x20 | 129,119 | 176.71x169.20x158.08 | 4,726,517.19 |
| | 2 | 20x20x20 | 106,110 | 157.91x167.75x149.72 | 3,965,840.17 |
| | 3 | 20x20x20 | 93,077 | 162.47x160.48x135.68 | 3,537,544.77 |
| | 4 | 20x20x20 | 123,177 | 186.45x175.47x137.99 | 4,514,651.81 |
| CENP-A | 1 | 20x20x20 | 101,693 | 168.22x160.48x141.59 | 3,822,262.68 |
| | 2 | 20x20x20 | 94,798 | 156.38x160.52x143.24 | 3,595,643.00 |
| | 3 | 20x20x20 | 147,710 | 174.34x174.34x174.34 | 5,298,474.34 |
| | 4 | 20x20x20 | 109,371 | 161.99x194.99x128.39 | 4,055,417.56 |
| CENP-$A_{Tail+Latch}$ $H3_{Core}$ | 1 | 20x20x20 | 101,774 | 171.05x178.99x125.31 | 3,836,636.98 |
| | 2 | 50x30x50 | 227,885 | 226.17x183.13x189.62 | 7,854,011.99 |
| | 3 | 50x30x50 | 236,237 | 223.99x196.90x184.86 | 8,152,726.70 |
| | 4 | 20x20x20 | 110,414 | 165.73x178.42x139.20 | 4,116,145.75 |
| $H3_{Tail+Latch}$ CENP-$A_{Core}$ | 1 | 20x20x20 | 150,769 | 175.78x175.78x175.78 | 5,431,682.75 |
| | 2 | 50x30x50 | 232,210 | 223.39x195.02x183.70 | 8,003,269.92 |
| | 3 | 20x20x20 | 128,166 | 173.80x164.98x163.47 | 4,687,329.87 |
| | 4 | 50x30x50 | 225,524 | 224.18x186.92x185.58 | 7,776,353.32 |
| Tailless H3 | 1 | 50x30x50 | 251,007 | 226.70x173.97x218.22 | 8,606,371.88 |
| | 2 | 50x30x50 | 225,997 | 223.99x176.55x196.09 | 7,754,249.02 |
| | 3 | 50x30x50 | 255,138 | 219.28x184.78x215.10 | 8,715,347.00 |
| | 4 | 50x30x50 | 255,678 | 238.14x185.88x197.37 | 8,736,629.01 |
| Tailless CENP-A | 1 | 50x30x50 | 277,074 | 234.72x194.56x206.74 | 9,441,091.91 |
| | 2 | 50x30x50 | 249,584 | 238.86x172.26x208.25 | 8,568,347.79 |
| | 3 | 50x30x50 | 251,035 | 226.70x192.33x197.77 | 8,622,888.81 |
| | 4 | 50x30x50 | 280,184 | 218.05x199.06x219.74 | 9,537,555.39 |

**Table S12. All-Atom Widom 601 DNA Post-Equilibration/Production Water Box Dimensions.** Water box solvation buffers, number of water molecules, and initial box dimensions for all-atom Widom 601 NCPs during production phase (2 μs). Solvation buffer is the specified distance. Initial box dimensions are the actual water box XYZ dimensions which encase the all-atom nucleosome structures.

| System | Run # | Solvation buffer (Å) | No. of water molecules | Initial box XYZ dimensions (Å) | Initial box volume ($Å^3$) |
|---|---|---|---|---|---|
| H3 | 1 | 50x30x50 | 239,827 | 225.76x179.36x204.00 | 8,260,416.23 |
| | 2 | 50x30x50 | 259,410 | 227.91x179.36x217.95 | 8,909,138.40 |
| | 3 | 50x30x50 | 233,544 | 233.36x186.23x185.41 | 8,057,636.85 |
| | 4 | 50x30x50 | 248,579 | 222.98x189.63x202.29 | 8,553,794.77 |
| CENP-A | 1 | 50x30x50 | 274,176 | 241.60x212.95x182.48 | 9,388,279.26 |
| | 2 | 50x30x50 | 262,233 | 225.12x192.44x206.93 | 8,964,903.20 |
| | 3 | 50x30x50 | 235,561 | 223.99x189.93x191.53 | 8,147,846.80 |
| | 4 | 50x30x50 | 234,298 | 215.24x182.41x206.74 | 8,116,475.16 |
| CENP-$A_{Tail+Latch}$ $H3_{Core}$ | 1 | 50x30x50 | 241,523 | 231.41x192.44x187.08 | 8,331,199.72 |
| | 2 | 50x30x50 | 265,985 | 218.05x197.84x210.98 | 9,101,537.16 |
| | 3 | 50x30x50 | 233,910 | 218.59x182.56x201.69 | 8,048,174.35 |
| | 4 | 50x30x50 | 235,144 | 221.11x191.30x191.84 | 8,114,223.25 |
| $H3_{Tail+Latch}$ CENP-$A_{Core}$ | 1 | 50x30x50 | 267,041 | 218.05x194.99x215.10 | 9,145,634.03 |
| | 2 | 50x30x50 | 333,004 | 261.72x197.47x217.95 | 11,264,028.24 |
| | 3 | 50x30x50 | 254,370 | 219.11x192.33x206.93 | 8,720,173.43 |
| | 4 | 50x30x50 | 271,228 | 223.39x191.30x217.05 | 9,275,370.16 |
| Tailless H3 | 1 | 50x30x50 | 241,868 | 223.99x187.84x197.37 | 8,304,202.26 |
| | 2 | 50x30x50 | 271,591 | 242.85x176.79x214.27 | 9,199,242.02 |
| | 3 | 50x30x50 | 236,251 | 225.12x184.78x195.01 | 8,112,114.54 |
| | 4 | 50x30x50 | 256,556 | 239.97x180.66x202.66 | 8,785,767.60 |
| Tailless CENP-A | 1 | 50x30x50 | 234,823 | 224.18x185.16x194.60 | 8,077,429.70 |
| | 2 | 50x30x50 | 237,092 | 232.30x176.52x199.65 | 8,186,460.80 |
| | 3 | 50x30x50 | 288,412 | 253.28x171.60x225.94 | 9,820,106.60 |
| | 4 | 50x30x50 | 246,763 | 220.24x183.13x209.59 | 8,453,467.40 |

**Table S13. All-Atom Alpha-Satellite DNA Post-Equilibration/Production Water Box Dimensions.** Water box solvation buffers, number of water molecules, and initial box dimensions for all-atom alpha-satellite NCPs during production phase (2 μs). Solvation buffer is the specified distance. Initial box dimensions are the actual water box XYZ dimensions which encase the all-atom nucleosome structures.
